# Comparative population genomics provide new insight into the evolutionary history and adaptive potential of World Ocean krill

**DOI:** 10.1101/2023.04.16.537052

**Authors:** Marvin Choquet, Felix Lenner, Arianna Cocco, Gaëlle Toullec, Erwan Corre, Jean-Yves Toullec, Andreas Wallberg

**Affiliations:** Department of Medical Biochemistry and Microbiology, Uppsala University; Husargatan 3, 751 23 Uppsala, Sweden; Natural History Museum, University of Oslo, Oslo, Norway; Department of Immunology, Genetics and Pathology, Uppsala University; Husargatan 3, 751 23 Uppsala, Sweden; Laboratory for Biological Geochemistry, School of Architecture, Civil and Environmental Engineering, École Polytechnique Fédérale de Lausanne (EPFL), 1015 Lausanne, Switzerland; CNRS, Sorbonne Université, FR 2424, ABiMS Platform, Station Biologique de Roscoff, Roscoff, France; Sorbonne Université, CNRS, UMR 7144, AD2M, Station Biologique de Roscoff, Roscoff, France

**Keywords:** krill, climate change, population genomics, comparative genomics, genetic adaptation, evolution

## Abstract

Genetic variation is instrumental for adaptation to new or changing environments but it is poorly understood how it is structured and contributes to adaptation in pelagic species without clear barriers to gene flow. Here we use extensive transcriptome datasets from 20 krill species collected across the Atlantic, Indian, Pacific and Southern Oceans and compare genetic variation both within and between species across thousands of genes. We resolve phylogenetic interrelationships and uncover genomic evidence in support of elevating the cryptic *Euphausia similis* var. *armata* into species. We estimate levels of genetic variation and rates of adaptive protein evolution among species and find that these are comparably low in large Southern Ocean species endemic to cold environments, including the Antarctic krill *Euphausia superba*, suggesting their adaptive potential to rapid climate change may also be low. We uncover hundreds of candidate loci with signatures of adaptive divergence between krill native to cold and warm waters and identify candidates for cold-adaptation that have also been detected in Antarctic fish, including genes that govern thermal reception such as *TrpA1*. Our results suggest parallel genetic responses to similar selection pressures across Antarctic taxa and provide new insights into the adaptive potential of important zooplankton that are already strongly affected by climate change.

## Introduction

The world’s oceans have absorbed over 90% of anthropogenic greenhouse gas energy released over the last century and warmed by about 1°C (Intergovernmental Panel on Climate Change 2014). This has strongly impacted pelagic species, causing rapid poleward shifts and accelerated phenologies in fish and zooplankton, such as jellyfish, salps, copepods and krill (Richardson 2008; Poloczanska et al. 2013; Poloczanska et al. 2016; Ratnarajah et al. 2023), and threatening to destabilize important food webs and ecosystem services (Doney et al. 2012; Baxter and Laffoley 2016). Genetic adaptation could be key to sustain marine populations, promoting resilience by targeting genes encoding traits such as growth, reproductive timing and thermal tolerance (Hoffmann and Sgrò 2011; Dam 2013). However, despite their pivotal roles in marine ecosystems, insight into what genes and variation that contribute to fitness and adaptation is still missing in zooplankton (Dam 2013; Bucklin et al. 2018).

Krill (Euphausiacea; “euphausiids”; 86 spp.) are crustacean macrozooplankton and grazers of phytoplankton primary production or smaller zooplankton (Russell et al. 1969). As important food for fish, mammals and birds, they play critical roles in transferring nutrients to higher trophic levels in marine ecosystems (Tarling 2010; McBride et al. 2014; Siegel 2016; Johnston et al. 2022) and superabundant species like the Antarctic krill *Euphausia superba* contribute to global biogeochemical systems through the biological carbon pump (Cavan et al. 2019). Different species occur throughout tropical, temperate and arctic ecosystems and their biogeography depend on physiological thermal tolerance, oceanographic conditions and nutrient availability (Russell et al. 1969; Cimino et al. 2020). Across these environments, euphausiids have evolved divergent life cycle strategies: low-latitude species associated with nutrient-rich upwellings or near-shore habitats tend to develop and mature quickly, be short-lived (<1 year or 1–2 years), breed multiple times or continuously throughout the year and have high productivity rates (Siegel 2000). High-latitude species instead deploy strategies to cope with extended periods of dark, cold and nutritionally adverse conditions, and are characterized by slow rates of development, extended longevity (e.g. 2 years or more), larger body sizes that build long-lasting lipid stores, and have short annual reproductive seasons and low productivity (Falk-Petersen et al. 2000; Siegel 2000; Siegel 2016). Moreover, many krill are stenothermal and occupy habitats with narrow latitudinal and thermal ranges. The Antarctic *E. superba* and *E. crystallorophias* likely split from other Southern Ocean species about 25–27 MYA, during the formation of the Antarctic Circumpolar Current (Zane and Patarnello 2000) and have since adapted to the extreme Antarctic environments. Today, *E. superba* inhabits cold waters ranging between −2.0°C to +4.0°C (Siegel 2016), while the neritic Ice krill *E. crystallorophias* occurs within an even more narrow thermal envelope (−1.8°C to 0°C) (Cuzin-Roudy et al. 2014). Short-term stress tests (e.g. CT50 assays) show they are sensitive to high temperatures (Cascella et al. 2015; Huenerlage et al. 2016), which is common among Antarctic ectotherms (Peck et al. 2014). In contrast, North Atlantic krill such as *Meganyctiphanes norvegica* (“the Northern krill”) and several *Thysanoessa* have thermal ranges spanning 2–15°C or more and occur from the colder Arctic Ocean to the warmer Gulf of Maine (Russell et al. 1969; Tarling 2010; Ollier et al. 2018), indicating greater thermal tolerance in these widely distributed euphausiids.

Similar to many other Antarctic species, *E. superba* and *E. crystallorophias* appear to have reduced capacity to upregulate inducible and protective heat shock proteins (Hsps) in response to increased temperature (Cascella et al. 2015; Huenerlage et al. 2016; Peck 2016; Chen et al. 2018; Toullec et al. 2020). Adaptation to cold environments could commonly involve fundamental genetic and physiological alterations and trade-offs that are maladaptive under rapidly warming temperatures (Pörtner et al. 2007), and it is therefore concerning that both Antarctic and Arctic krill are declining or shifting to higher latitudes due to climate change (Atkinson et al. 2019; Edwards et al. 2021). Tropical krill may also live close to their thermal limits and be sensitive to change, as suggested in other marine invertebrates (Peck et al. 2014), although these species are not regularly monitored or studied. The long-term ability of krill, or other planktonic organisms, to resist temperature variation is difficult to estimate experimentally and it is not clear if any krill has enough physiological tolerance to track continued ocean warming.

Evolutionary adaptation through natural selection on adaptive variation could be crucial for coping with climate change. Knowledge about the genetic mechanisms that contribute to adaptation can be gained through genome-scale comparisons between populations or species adapted to different local conditions (Savolainen et al. 2013; Meek et al. 2023). In marine taxa, scans for adaptive divergence between populations have uncovered loci involved in reproductive timing in Atlantic herring (Han et al. 2020) and bleaching in *Acropora* corals (Fuller et al. 2020), while comparative analyses among *Mytilus* mussel species have detected candidate genes evolving under divergent temperature-dependent selection (Popovic and Riginos 2020). Population genetic theory suggests that zooplankton could have high adaptive potential (Peijnenburg and Goetze 2013): i) they typically have large populations with many reproductive individuals (i.e. a high effective population size, or “*N*_e_”) that can maintain or generate much variation to select from; ii) due to large *N*_e_, they are expected to be comparably unaffected by genetic drift, i.e. stochastic changes in allele frequencies between generations that may interfere with selection; iii) many species have short generation times and high reproductive rates, amenable for adapting to rapid changes. However, empirical support for these predictions is still missing as various challenges have hindered molecular research on marine zooplankton.

One of these challenges resides in the ambiguity of taxonomy across zooplankton taxa (Bucklin et al. 2007; Bailey et al. 2016; Choquet et al. 2018). Thus, the widespread occurrence of similar-looking or cryptic species has led to under-estimation of zooplankton diversity, although large-scale barcoding and metabarcoding surveys have started to address this issue (Bucklin et al. 2016; Bucklin et al. 2021). In krill, cryptic variation was reported within *Stylocheiron affine* (Wiebe et al. 2016) and the specific status of *Euphausia similis* and its variety *armata* (Hansen 1911) remain unclear. Furthermore, accessing zooplankton species potentially adapted to polar environments can be a logistical challenge when they live in remote locations, like the Antarctic krill, and may therefore discourage attempts at conducting advanced studies of local adaptation (Bucklin et al. 2018). Besides, such studies require genome-wide information and yet only a handful of genomes have been sequenced so far across zooplankton, mostly due to very large genome sizes common in many marine zooplankton, which, combined with a small body size, prohibit whole genome sequencing (Bucklin et al. 2018). Krill have huge and repetitive genomes that, among six known species, range between 11– 48 Gb (Jeffery 2012), making genome-wide analyses very challenging. To date, only the 48 Gb genome of the Antarctic krill has been assembled and population-scale scans recovered few potentially adaptive variants (Shao et al. 2023), possibly due to the highly panmictic life history of this species (Bortolotto et al. 2011; Deagle et al. 2015; Shao et al. 2023). Overcoming these obstacles is critical for monitoring and forecasting zooplankton responses to climate change. While the constant advances of sequencing technologies and protocols promise to ease whole genome sequencing of challenging organisms in the future, transcriptomics offers many possibilities to advance insight into zooplankton ecology (Lenz et al. 2021).

Here we produce, for the first time, a comprehensive comparative dataset at the genome-scale for krill, spanning 20 species from the Atlantic, Indian and Southern Oceans (fig. 1). Euphausiids are an attractive model for zooplankton because of their importance for ocean ecosystems and distribution across all latitudes. Moreover, they belong to a comparatively small and ecologically well-characterized order of zooplankton and are not extremely divergent, facilitating association between genes and phenotypes through comparative analyses. We sidestep the issue of prohibitively large genomes of these non-model organisms by using RNA-seq data to analyze genetic variation in expressed protein-coding genes (De Wit et al. 2015). We aim to elucidate the evolutionary history of krill by comparing variation between species, uncovering candidate genes for adaptation to cold or warm environments and testing if krill have high adaptive potential, as well as identify factors that may determine it. In addition, we provide further evidence for how thermal tolerance varies between species and associate with the environment.

**Fig. 1.**
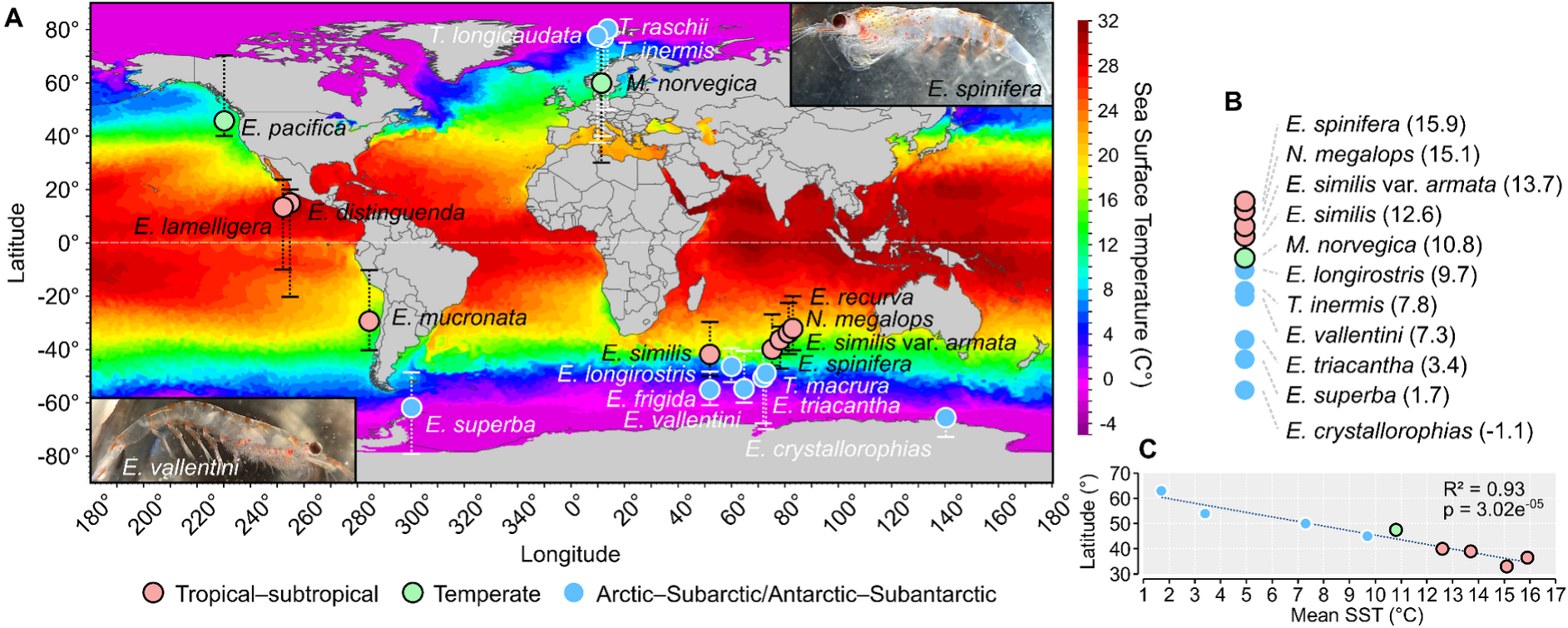
The geographic distribution of 20 surveyed krill species. (*A*) Circles indicate locations of one or more collected specimens (colors represent the thermal conditions associated with the range of each species). The basin-specific latitudinal range of each species is indicated with bars. Ranges are from Russel & Yonge (Russell et al. 1969). Global Sea Surface Temperature readouts are a daily snapshot from OISST V2 (2022-06-01) from ERDDAP: https://www.ncei.noaa.gov/erddap/griddap/. Photos of Indian Ocean krill collected in 2019. (*B*) Approximate mean SST of the ranges of 11 species used for analyses of thermal tolerance or intraspecific variation. (*C*) Linear regression correlation between habitat latitudinal midpoint and mean SST (n=9 species).

## Results

### Thermal tolerance in krill is associated with habitat temperature

We used CT50 mobility assays to characterize and compare thermal tolerance in six krill species sampled from the Southern, Atlantic and Indian Oceans. The two Antarctic species—the Ice krill (*E. crystallorophias*) and the Antarctic krill (*E. superba*)—have significantly lower CT50 thresholds compared to the two subantarctic *E. vallentini* and *E. triacantha* (fig. 2*A*). The average CT50 for *E. vallentini* is 17.9±0.42°C. This may be an underestimate because specimens were fragile and appeared impacted by fishing (reflected in the rapidly subsiding high sigmoid plateau and the flatter slope). CT50 for *E. triacantha* was 18.37±0.39, which appeared to be at better health. The widespread Northern krill *Meganyctiphanes norvegica* showed a record CT50 with a value of nearly 23°C for animals that were fished in 4°C waters of the high Arctic Spitsbergen (same locality as *T. inermis*), clearly illustrating extreme tolerance to temperature in a species ranging from the warm Mediterranean Sea to cold Arctic waters (Tarling 2010). CT50s are strongly correlated with ambient temperatures (fig. 2*B*; supplementary fig. S1), underscoring that thermal tolerances of krill are linked to the respective temperatures of their native environments.

**Fig. 2.**
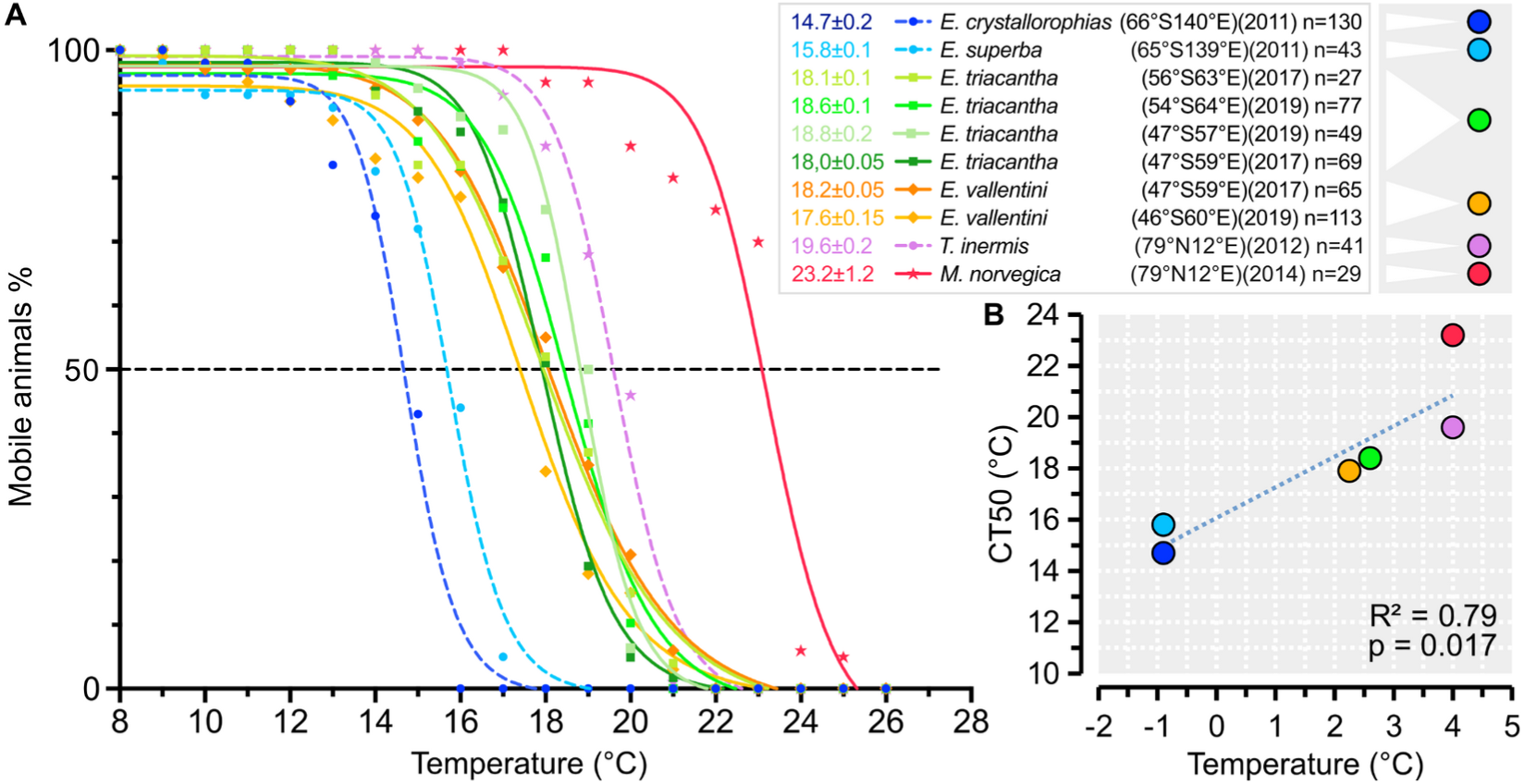
Thermal tolerance in Southern Ocean and Atlantic Ocean krill. (*A*) The figure shows all the results acquired over several years during campaigns in Antarctica, Arctic and Indian Ocean. Three species (dashed curves) have been published before (Cascella et al. 2015; Huenerlage et al. 2016), while *E. vallentini* (orange), *E. triacantha* (green) and *M. norvegica* (red) are new to this study. The white legend indicates species/experiment-specific calculated CT50 values, fishing coordinates, years and sample sizes. (*B*) Linear regression correlation between CT50 and the measured ambient sea temperature at the experimental locations. The mean CT50 of all experiments was used for *E. triacantha* (n=4) and *E. vallentini* (n=2).

### Genome-scale datasets to study variation and adaptation

We compiled RNA-seq data spanning 124 transcriptomes from 20 species (102 transcriptomes from 17 species are new for this study). Morphological and molecular species identifications agreed for all samples with barcodes in the MetaZooGene database (Table S1). After removing redundant and non-coding isoforms, the reference transcriptomes contained 20–53 K coding transcripts per species (Table S2). BUSCO analyses indicated high completeness (median=92.5%) and low duplication levels (median=3.3%), amenable for tracing evolutionary patterns using base-level substitutions and polymorphisms. We measured intraspecific genetic variation in nine species, using up to 20 specimens per species and detected between 260–2,200 K high-quality SNPs across 12–22 K genes (Table S3).

### Genetic divergence between Indian Ocean *Euphausia similis* and *Euphausia similis* var. *armata* suggests they are different evolutionary lineages

The Indian Ocean sampling region spanned multiple waterfronts and we interrogated samples from this region for population structure. Two out of three temperate–subtropical species showed structure. We sequenced 19 specimens of *E. similis* from the Indian Ocean (fig. 3*A*).

**Fig. 3.**
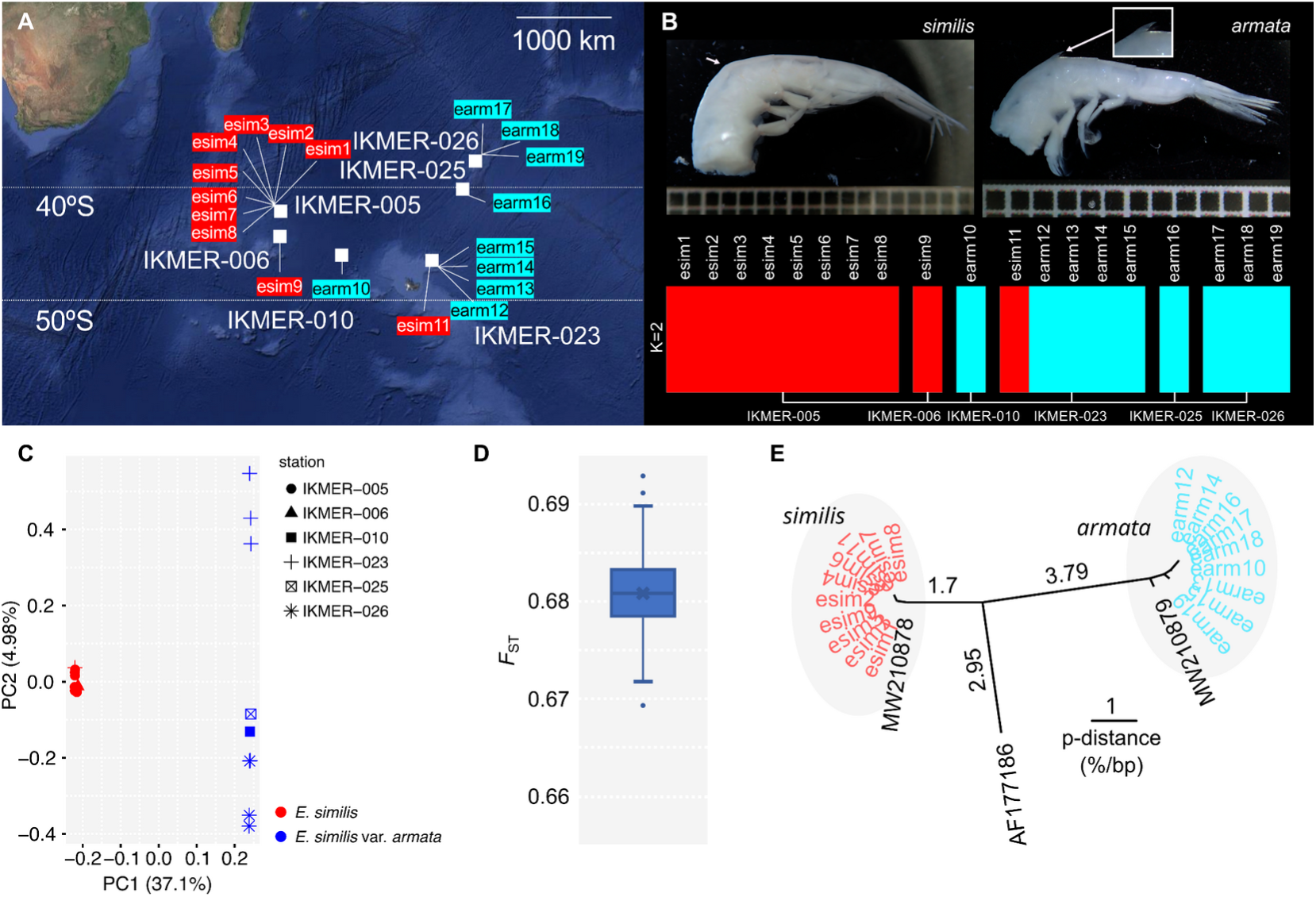
Genetic structure between *Euphausia similis* (“esim”) and *Euphausia similis* var. *armata* (“earm”) from the Indian Ocean. (*A*) Map of sampling stations (n=6). (*B*) Top: representative tails of *E. similis* (without a dorsal spine) and *E. similis* var. *armata* (with a spine on the third abdominal segment). Arrows indicate location of the spine (if present). Bottom: genetic ancestry and admixture of samples (K=2 ancestral groups). (*C*) The genetic structure among 19 samples using PCA (n=84,436 unlinked nuclear SNPs). (*D*) Weighted *F*_ST_ estimated from 1,000 random resamples drawing 1 SNP per gene (n=6,074 genes) (Choquet et al. 2019). (*E*) A mitochondrial COI neighbor-joining gene tree inferred from uncorrected pairwise genetic distances. Distances are shown for the major internal branches (% differences per bp). Sequences from MetaZooGene are: AF177186 (north-east of Japan; Kenji Taki; personal communication); MW210878 and MW210879 (the southern Atlantic Ocean).

Nine specimens collected mostly to the east morphologically matched the *Euphausia similis* var. *armata* variety (“*armata*”), having an accessory abdominal spine of variable size (fig. 3*B*; supplementary fig. S2*A*) (Hansen 1911; Baker et al. 1990). *E. similis* without the spines were mostly found to the west (figs. 3*A*–*B*; supplementary fig. S2*B*; “*similis*”). We initially detected strong structuring between *similis* and *armata* using ancestry analysis and PCA (fig. 3*B*–*C*), which was also supported by high *F*_ST_ (0.68) (fig. 3*D*). We next assembled 500–800 bp mtCOI fragments and assessed divergence. Sequences segregated consistently with the nuclear data and were ∼6.2% divergent, clustering with different MetaZooGene *similis* barcodes derived from southern Atlantic Ocean samples (fig. 3*E*; supplementary fig. S3). Because of the consistent molecular and phenotypic differences between *similis* and *armata*, we treated them as separate evolutionary lineages in subsequent analyses. We detected comparably weak north–south structuring in our limited *N. megalops* sample (*F*_ST_=0.06), but did not observe any structure in the subtropical *E. spinifera*, nor in the subantarctic *E. vallentini* and *E. triacantha* collected from the Antarctic Polar Front (supplementary fig. S4).

### Two-fold difference in genetic diversity among nine species of krill

We took transcriptome-wide levels of genetic variation as indicators of genetic diversity and studied how it varied among gene regions and species (supplementary figs. S5–S6). Overall, we detected about 1.6× more variation in untranslated regions (UTRs) compared to coding regions (average *θ*_w_=1.4% vs. 0.9%), and 3.1–5.6× more variation at synonymous sites compared to non-synonymous sites (fig. 4; supplementary fig. S5). These patterns likely reflect both direct and linked purifying selection, reducing variation around functionally important sites (Cvijović et al. 2018). Restricting measurements to only genes with established orthology (see below), we observe lower but strongly correlated levels of variation (∼0.78×; r^2^=0.99; supplementary fig. S7), indicating the full gene sets are less conserved than orthologs overall but representative of variation among species. We let synonymous variants represent selectively neutral variation and be indicative of effective population size (*N*_e_) and demographic history. All species have variation surpassing 1% per-base, the most diverse being *E. vallentini* (*π*_S_=2.5%; *θ*_w_=3.83%), while *E. triacantha* is the least diverse (*π*_S_=1.1%) (fig. 4). Assuming similar mutation rates among species, estimates of *N_e_* indicate *E. vallentini* (3.6 M) and *E. similis* var. *armata* (3.2 M) have the largest effective population sizes (Table 1). Our *armata* sample has 1.7× as much variation as *similis sensu stricto*, further suggesting these lineages follow different evolutionary trajectories. Tajima’s D (D_T_) is negative across all nine species, ranging from −1.2 to −1.9 (Table 1), indicating excess low-frequency variants compared to expectations under neutrality, which could result from processes such as selective sweeps or recent population expansions. Neither intrinsic life history traits (e.g. body or larval size) or environmental parameters (e.g. temperature, latitude or range) are strong predictors of *π*_S_ or D_T_ (i.e. no correlation p<0.1; see Table S9 for parameters).

**Fig. 4.**
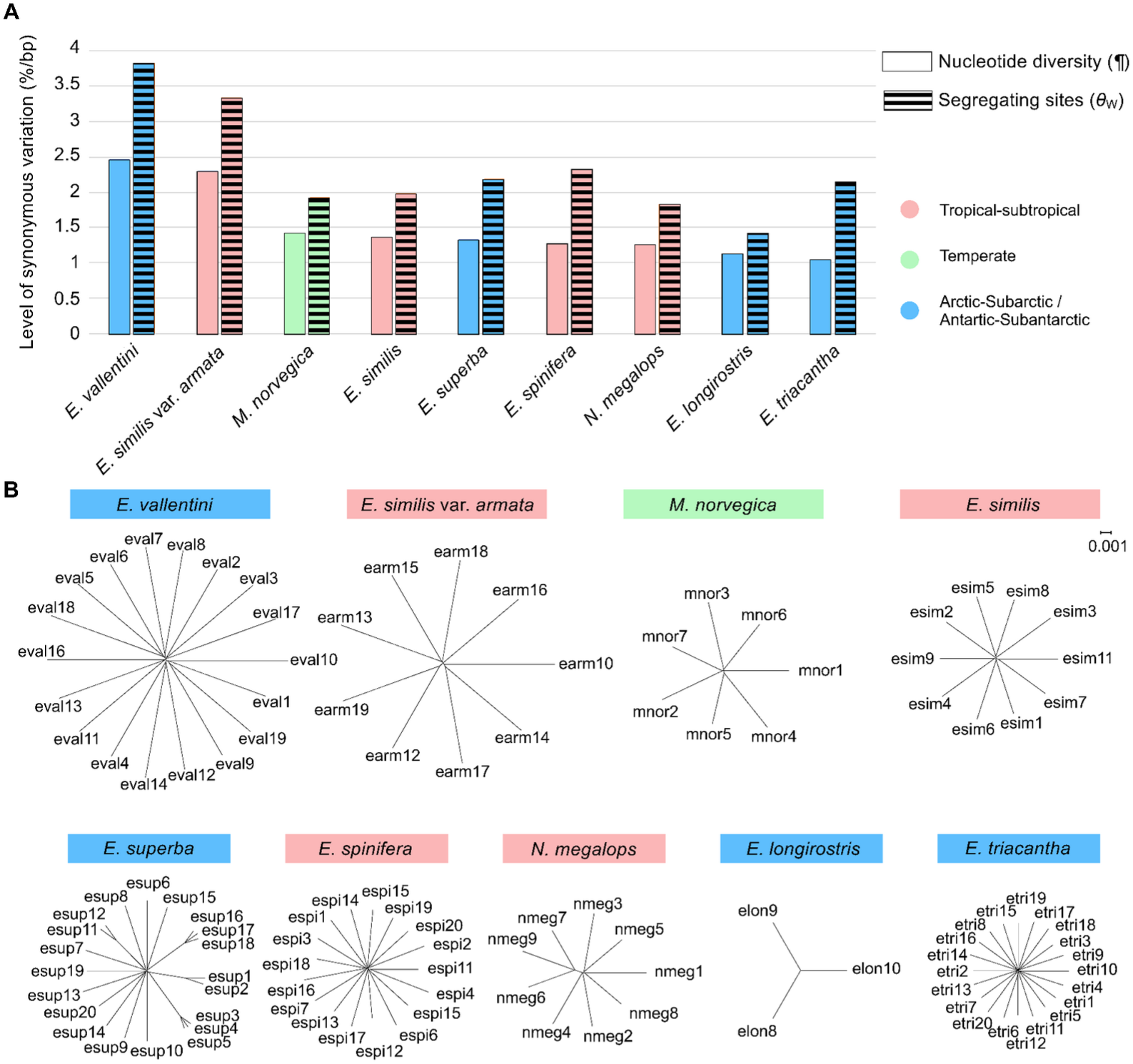
Genetic variation at synonymous sites in nine krill species. (*A*) Nucleotide diversity (*π*_S_) and Watterson’s theta (*θ*_wS_) estimated from 74 k to 709 k synonymous SNPs per species, after correction for numbers of accessible sites. (*B*) Neighbour-joining trees showing variation and structure across species. Branch lengths are scaled as the average genetic distance per base between samples (*π*_S_), estimated at synonymous loci and corrected for accessible sites and drawn at the same scale.

**TABLE 1.**
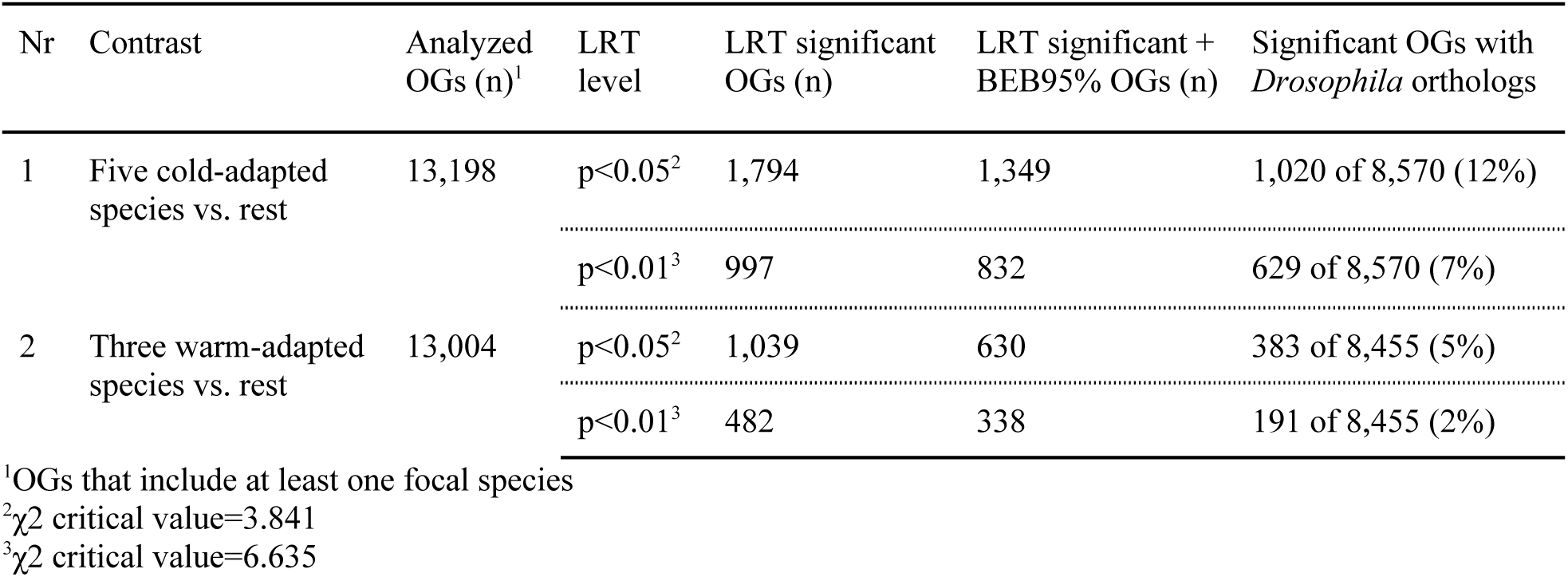
Synonymous variation and demographics in nine krill species.

### Comparative analyses establish comprehensive gene orthologies and a robust species tree

To enable direct comparative analyses of molecular evolution, we inferred gene orthology among the 20 krill species and seven outgroups (supplementary materials) and reconstructed a species tree. We detected 103,292 orthogroups (OGs) with genes from two or more krill species, but only 880 OGs contained single-copy orthologs spanning all 20 species. We produced two subsets of OGs, balancing species missingness against the number of OGs. The first dataset spanned 2,280 OGs, >1 million amino acid positions and 18+ krill species (Table S4), and was used to reconstruct a fully supported phylogeny (fig. 5*A*). The topology is mostly consistent with inferences of krill interrelationships by Vereshchaka and co-workers based on morphology and four molecular markers: *Meganyctiphanes* + *Nematoscelis* + *Thysanoessa* form the monophyletic clade *Nematoscelinae*, which is the sister taxon of *Euphausiinae* (Vereshchaka et al. 2019). Groupings within *Euphausia* confirm a monophyletic Southern Ocean “*Euphausia superba* group” including *frigida*, *vallentini*, *crystallorophias* and *superba*, while our clade with *spinifera*, *longirostris*, *similis*, *similis* var. *armata* and *triacantha* also include *recurva* and *mucronata*, mixing other species groups.

**Fig. 5.**
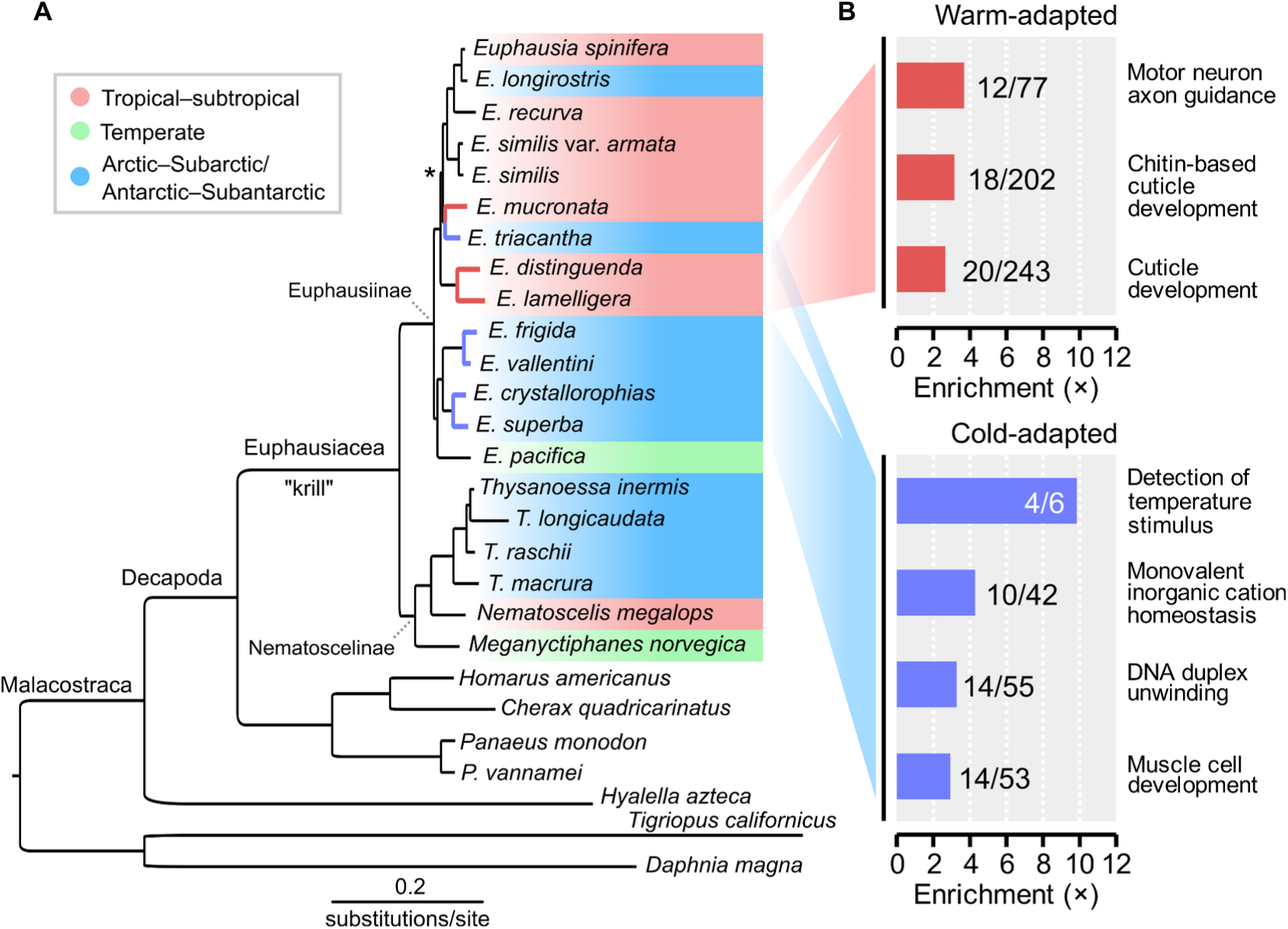
Phylogenomic inference of interrelationships and molecular evolution in krill. (*A*) A species tree inferred from 2,280 orthologous protein alignments and constructed from 1,000 ultrafast bootstrap replicates (JTTDCMut+F+I+G4 model; lnL=-11,173,074) (Hoang et al. 2018). All nodes in the majority-rule consensus tree but one have 100% bootstrap support (*= 99% support). Foreground branches used in branch-site tests to detect positively selected genes are highlighted (blue=five cold-adapted species; red=three warm-adapted species). (*B*) Statistically enriched gene ontologies (GO) among candidate genes for positive selection (False Discovery Rate <0.05) in the “cold-adapted contrast” (n=629 candidates; LRT<0.01 with ≥1 BEB95% sites) and “warm-adapted” ontologies (n=383 candidates; LRT<0.05 with ≥1 BEB95% sites). Numbers beside bars indicate genes in the target set (i.e. selection candidates) vs. genes in the background test set. Redundant GO terms were removed with *Revigo* (Supek et al. 2011).

Resolving phylogenetic interrelationships is necessary for understanding the evolution of euphausiid genes, traits and ecologies and numerous attempts have been made using morphology or sparse molecular datasets such as mitochondrial markers (see Bucklin et al. 2007; Vereshchaka et al. 2019; Schram and Koenemann 2022 for reviews). Our reconstruction is the first phylogenomic species tree for krill and shows that *E. triacantha* is only distantly related to other *Euphausia* of the Southern Ocean. *E. triacantha* and *E. longirostris* have likely expanded into cold environments independently from each other, and from other *Euphausia*. Our tree implies that krill have frequently diversified into different climates, offering the opportunity for studying parallel events to uncover genetic adaptations.

### Scans for positive selection reveal candidate genes for environmental adaptation

We aimed to detect genes that may underlie episodic adaptation to cold or warm climates in krill. To this end, we applied branch-site tests to the OG dataset spanning >13,000 OGs (>5 million codon positions; Table 2; Table S4) to detect genes with locally elevated ω between species associated with different environments (Yang and Nielsen 2002). In the first contrast, we compared five Southern Ocean cold-adapted *Euphausia* against the rest of the phylogeny (*E. superba*, *E. crystallorophias*, *E. frigida*, *E. vallentini* and *E. triacantha*; fig. 5*A*; Table S4). We identified 832 gene candidates (LRT p<0.01; n=629 genes with *Drosophila* homologs; Table S5), containing 1,800 selected codons. A gene ontology (GO) enrichment test found enrichment for genes involved in detecting temperature among candidates (fig. 5*B*; Table S6), including homologs of *Transient receptor potential cation channel A1* (*TrpA1*), *painless*, *subdued* and *straightjacket*. Both *TrpA1* and *painless* encode TRPA ion channels (Hwang et al. 2012; Himmel and Cox 2020), which are activated by temperature and various ligands. *TrpA1* is linked to sensation of either harmfully cold or warm temperatures across the animal tree of life (Zhang et al. 2022), while *painless* is associated with heat sensation (Dhaka et al. 2006; Himmel and Cox 2020). Crustacean TRP evolution is characterized by widespread gene duplication (Kozma et al. 2020), and phylogenetic analysis reveals multiple paralogs also in krill, and suggest the krill *TrpA1* OG is orthologous to decapod *TrpA1-like*, while the *painless* OG could be a “*Pain1*” ortholog (supplementary fig. S8). We detect one intracellular and one extracellular site with significant evidence of positive selection in cold-adapted species in the otherwise conserved *TrpA1* OG (ω=0.171; fig. 6). The *painless* OG contains six significant sites but this gene is much more rapidly evolving overall (ω=0.392), which could indicate erroneous clustering of divergent paralogs in the orthology inference. *subdued* encodes an anoctamin ion channel of the Transmembrane protein 16 family (Yuan et al. 2022), which conducts chloride in response to heat and is associated with thermal nociception in *Drosophila* (Jang et al. 2015). It is conserved overall (ω=0.12) and we detect three sites with signatures of positive selection (supplementary fig. S9*A*). In addition, the candidate set contains six heat shock protein or chaperone genes, including homologs of *Hsp110*/*Hsc70Cb*, *Hsp67Bc*, *Hsp70Bbb* and three *Chaperonin containing TCP1* chaperonins (*CCT3*, *CCT7* and *CCT8*), that are involved in folding proteins and protecting them from noxious temperatures and other environmental stressors (Mayer and Bukau 2005; Chen et al. 2018).

**Fig. 6.**
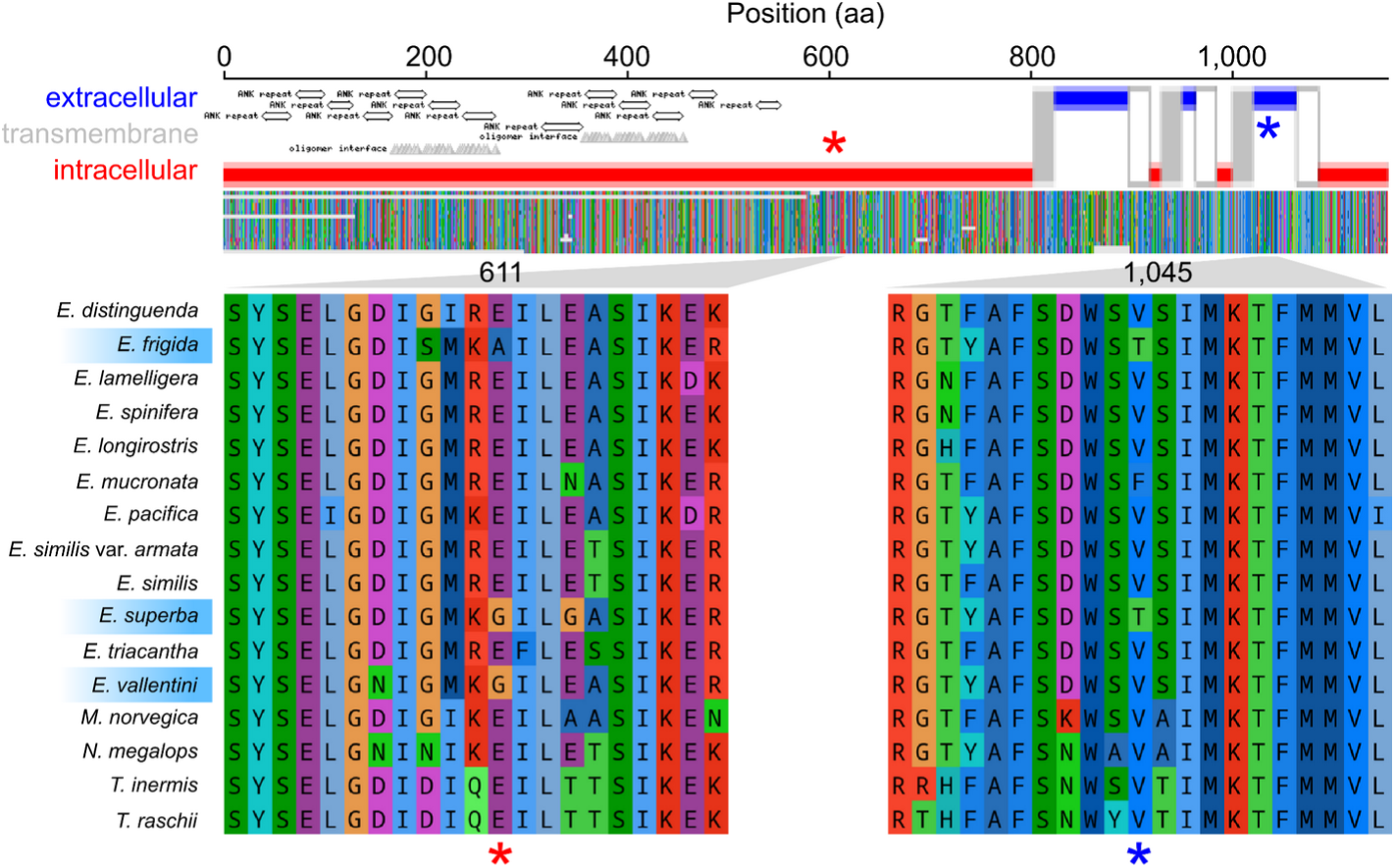
Signatures of positive selection in a *Transient receptor potential cation channel A1* (*TrpA1*) homolog in cold-adapted krill. Top: the intracellular, extracellular and six expected transmembrane domains of the TRPA1 membrane protein encoded by the *E. superba* ortholog (predicted using *TOPCONS v2.0* (Tsirigos et al. 2015)). Gray transmembrane regions=outbound; white=inbound. Ankyrin-repeats were detected using BLAST Conserved Domain Search (Lu et al. 2020). Middle: the trimmed protein multiple-sequence alignment (MSA) of the krill TRPA1 orthogroup. Bottom: magnified view of the MSA highlighting the sites and species. Two stars indicate putatively selected sites (BEB>95%).

**TABLE 2.**
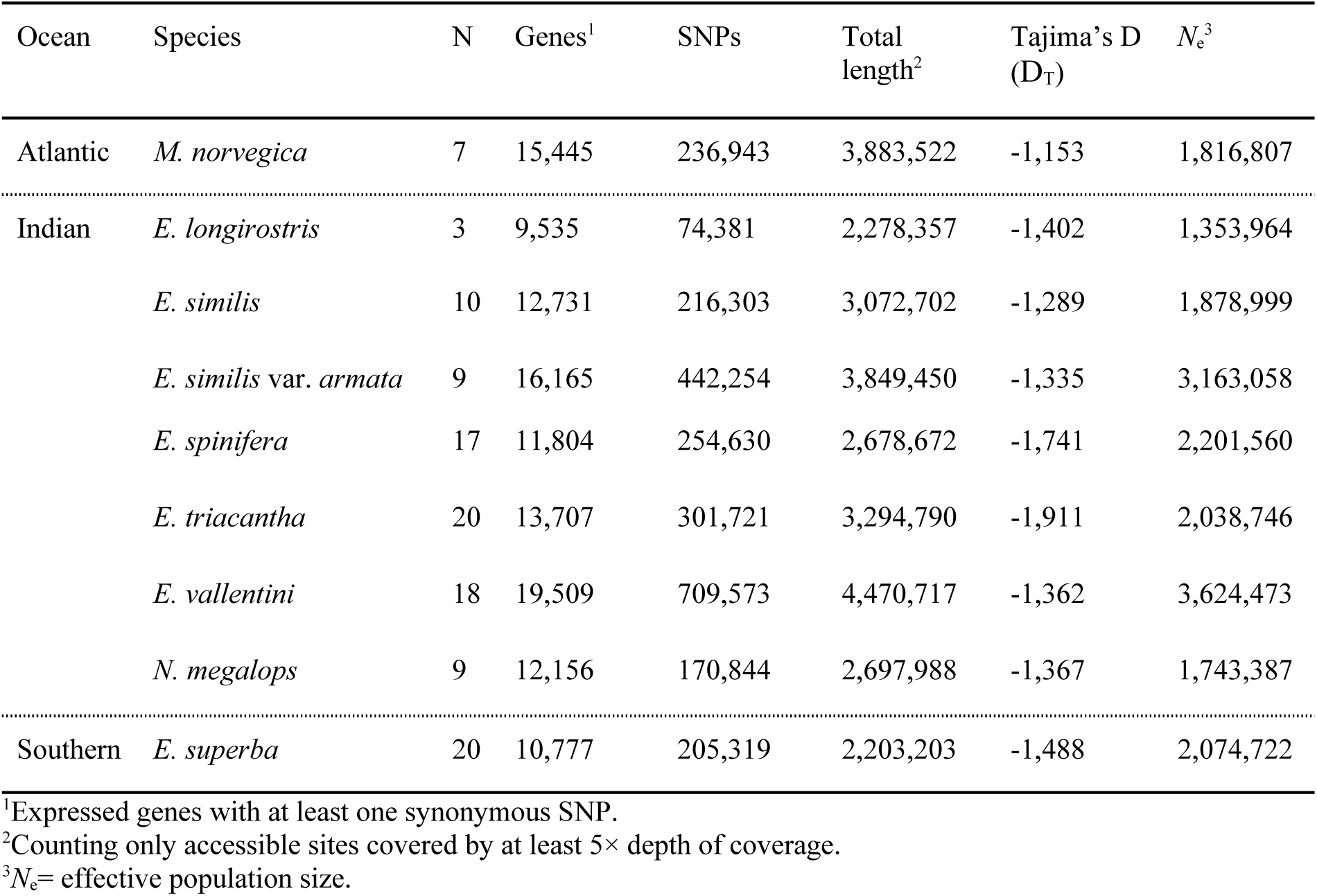
Signatures of positive selection on protein-coding genes.

In the second contrast, we compared three tropical–subtropical Pacific Ocean species against the rest ((*E. distinguenda*, *E. lamelligera* and *E. mucronata*; fig. 5*A*; Table S4). In this smaller foreground, we detected fewer candidates and less strong GO enrichments (383 genes with LRT<0.05). We found signatures of positive selection in genes involved in axon guidance and neurodevelopment (fig. 5*B*; Tables S5–S6), including *Neurotrophin 1* (*NT1*) and *Semaphorin 1a* (*Sema1a*) (Cafferty et al. 2006; Shen et al. 2017). In addition, we detect selection signals in two heat shock protein-related genes: *Hsc/Hsp70-interacting protein* (*HIP*) and *Heat shock protein 70 cognate 5* (*Hsc70-5*), the latter of which is linked to heat response in crustaceans (Yuan et al. 2017). This contrast uncovers several candidates for photoreception, including *ninaB*, which synthesizes the retinal chromophore pigments of rhodopsin (Voolstra et al. 2010) and *Photoreceptor dehydrogenase* (*Pdh*), which also processes retinal pigment. We identify an opsin (OG98713; supplementary fig. S9*B*), automatically annotated as Rh5. However, queries against KrillDB^2^ (Urso et al. 2022) and NCBI Genbank suggest it is a peropsin, a light-sensitive yet non-visual receptor with poorly understood function in crustaceans but with expression patterns that oscillate daily in Antarctic krill (Henze and Oakley 2015; Biscontin et al. 2016; Palecanda et al. 2022).

In both contrasts, we detect selection signals across multiple genes encoding cuticular proteins (n_cold_=10; n_warm_=5) and chitinases (n_cold_=1; n_warm_=1). Cuticular protein genes are among the largest gene families in arthropod genomes (Magkrioti et al. 2004), and help build the exoskeleton. In krill, cuticular development and gene expression are closely linked to physiological and seasonal cycles of ecdysis (molting), growth and reproduction (Buchholz and Buchholz 2010; Urso et al. 2022). In krill native to warm waters, we detect potentially selective substitutions in *Ecdysone-induced protein 75B* (*Eip75B*), a gene encoding a nuclear receptor that regulates the timing of molting, reproduction and circadian oscillation in insects and crustaceans (Kumar et al. 2014; Gouveia et al. 2018), that could contribute to adaptation.

### Slow rates of adaptive protein evolution in some Southern Ocean species

Theory suggests high efficacy of natural selection and adaptive potential in zooplankton (Peijnenburg and Goetze 2013). This hypothesis predicts that much variation between related zooplankton should have evolved through adaptation. To test this, we estimated how much of divergence in protein-coding genes between species may have been driven by positive selection as opposed to non-adaptive processes (like genetic drift) in krill. We inferred *dN* and *dS* between species (Tables S7–S8) and ancestral nodes and estimated intraspecific polymorphism (*pN/pS*) using unfolded site frequency spectra (supplementary fig. S10). We next estimated the rates of adaptive and non-adaptive amino acid substitutions (ω_A_ and ω_NA_), respectively, and the overall proportions of adaptive amino acid substitutions (α) using non-parametric and model-based approaches (Table S7). We used the Gamma-Zero model to compare all species, which fits a gamma distribution for neutral and deleterious mutations to the data. It produced estimates similar to the Displaced-Gamma model, both of which behaved consistently across all datasets, producing estimates correlated with but higher than basic non-parametric methods and were considered accurate in recent tests (Al-Saffar and Hahn 2022). We detected the highest rates of adaptive protein evolution in the subtropical *E. similis* var. *armata* lineage (α=0.59; ω_A_=0.086) (fig. 7*A*–*C*; Table S8), whereas adaptive evolution appears more limited in the Antarctic krill *E. superba* (α=0.20; ω_A_=0.035) and subantarctic *E. triacantha* (α=0.30; ω_A_= 0.059). We then analyzed how effective population size (*N*_e_), here approximated by *π*_S_, life history traits and habitat conditions correlate with adaptive rates to identify factors that may influence adaptive potential. Since inference of α depends on *π*_S_, we used estimates of *π*_S_ from genes not used to compute α. We find that *π*_S_ predicts α (r^2^=0.53; p=0.040), but is more strongly associated with ω_NA_ (r^2^=0.54; p=0.039) than with ω_A_ (r^2^=0.37; p=0.108) (fig. 7*D*). Our results suggest that genetic variation between krill species may be more strongly driven by increased fixation of non-adaptive variants in small populations than by accelerated positive selection in large populations, a pattern previously observed across animal phyla (Galtier 2016). We find that small body and propagule sizes (i.e. larval sizes) and warmer habitats are strongly associated with high α (fig. 7*E*–*F*; Table S9), suggesting that life history and habitat characteristics may influence the amount and rates of adaptation in krill.

**Fig. 7.**
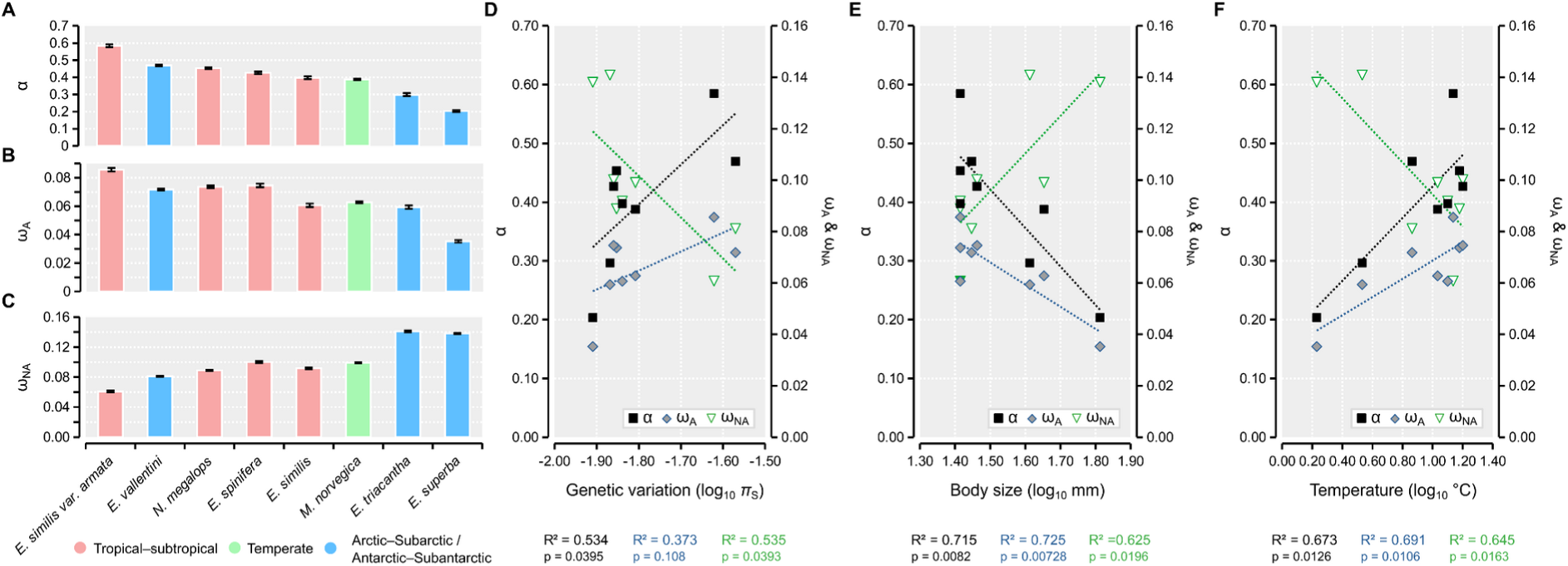
Estimates of adaptive protein evolution in nine krill species (GammaZero model). (*A–C*) The proportions and rates of adaptive and non-adaptive amino acid substitutions (α, ω_A_ and ω_NA_), respectively. Whiskers indicate maximum likelihood confidence intervals. (*D–F*) Log_10_ linear regression correlations between genetic variation (π_S_), maximum body size and habitat sea-surface temperature and α, ω_A_ and ω_NA_, respectively. R^2^ Pearson correlation coefficients and significance-values shown below the graph.

## Discussion

We here use comparative population genomics to characterize the patterns of genetic variation and evolutionary history of krill on the largest scale to date. Assessments of phylogenetic interrelationships and genetic adaptation among krill species have previously only been based on a few markers. Our analyses of molecular evolution in over 13,000 putative gene orthologs among krill native to cold and warm environments offer new insights into genetic variation, taxonomic diversity and adaptation.

### Elusive determinants of genetic variation in krill

Mitochondrial markers have uncovered considerable variability in intraspecific variation among krill species but rarely exceeding 2%/bp (Bucklin et al. 2007; Bortolotto et al. 2011). We here estimate levels of selectively neutral genetic variation across thousands of nuclear genes and find that it varies more than two-fold among nine species (*π*_S_=1.1–2.5%). Large and superabundant species like the Antarctic krill *E. superba* and Northern krill *M. norvegica*, that each have enormous biomass estimated to 100s of Mts (Atkinson et al. 2009; Tarling 2010), only have intermediate levels of variation. The most diverse species in our set is *E. vallentini* (*π*_S_=2.5%), a small (13–28mm) subantarctic, omnivorous and possibly panmictic species with a 2-year lifespan and circumpolar distribution mostly distributed north of the Polar Front (Russell et al. 1969; Ridoux 1988; Mayzaud et al. 2003; Harkins et al. 2013; Cuzin-Roudy et al. 2014). It can be the dominant species in shelf areas, including the Patagonian coast and the Kerguelen archipelago (where we collected it) (Palma and Silva 2004; Koubbi et al. 2011; González et al. 2016). In contrast, we find that *E. triacantha* is the least diverse species (*π*_S_=1.1%). This large (24–41mm) and carnivorous krill has a 3-year lifespan and 2-year generation time and a circumpolar distribution that spans the Polar Front, across which it can dominate the mesopelagic euphausiid biomass (Russell et al. 1969; Siegel 1987; Phleger et al. 2002; Cuzin-Roudy et al. 2014). These two extremes could represent “r-strategist” vs. “K-strategist” species, whose characteristic life history traits (e.g. short vs. long lifespan, small vs. large bodies or propagules) have previously been linked to high vs. low genetic diversity across metazoans (Romiguier et al. 2014; Ellegren and Galtier 2016). However, associated traits like fecundity are not known in either species, and across our dataset we otherwise find no statistically significant correlations between such life history traits and levels of variation. We are therefore unable to identify general determinants of diversity in krill, which may need sampling of more species and ecological information.

Compared to the neutral levels of genetic diversity of other arthropods, we find that krill has intermediary levels of variation (*π*_S_: mean±SD): 1.5±0.5% in krill vs. 1.4±0.9% in 43 species (Leffler et al. 2012) or 1.4%±1.2% in 26 species (Romiguier et al. 2014). Thus, krill are in this sense neither hypo- or hypervariable. Across the tree of life, genetic diversity in natural populations varies much less than census population sizes (*N*_c_), a largely unexplained phenomenon termed “Lewontin’s paradox” (Lewontin 1974; Charlesworth and Jensen 2022), and many marine species appear to have much lower *N*_e_ than expected from abundance (Hedgecock 1994). We infer long-term *N*_e_ for *E. superba* and *M. norvegica* to be only 2 M and 1.8 M, respectively, far below expected *N*_c_ of hundreds of trillions of individuals. Low estimates of *N*_e_ inferred from limited observable variation could be due to selection for low mutation rates afforded in species with large populations (Sung et al. 2012). For example, mutation accumulation lines in the ubiquitous unicellular phytoplankton *Emiliania huxleyi* recovered a low mutation rate (µ=5.6×10^−10^), although it was not considered low enough to explain its modest levels of variation (*π*_S_=0.6%) (Krasovec et al. 2020). Likewise, Shao and co-workers recently estimated low nucleotide diversity in the Antarctic krill (π_genome_≈0.25%; π_CDS_≈0.17%) using genome-scale data, and also inferred a low mutation rate (μ=6.2×10^−10^) as a means to explain this (Shao et al. 2023). These estimates are extremely low and difficult to reconcile with ours. We measure 5.3× as much variation at synonymous sites than the genome-wide estimate in *E. superba* (1.3% vs. 0.25%), or in coding regions overall (0.88% vs. 0.17%). Insight into genomic variation in euphausiids is still limited, as is baseline data derived from other sources than next-generation sequencing. However, Papot and co-workers studied variation in three PCR-amplified paralogs of the nuclear *hsp70* in *E. superba* and *E. crystallorophias* (Papot et al. 2016), and re-analysis of their *E. superba* data gives *π*_S_ ranging between 1.4–5.0% (π_S(A)_=1.4%; π_S(B)_=5.0%; π_S(C)_=1.8%), close to our average estimate across 10,777 genes. Moreover, Shao and co-workers derived low μ assuming a molecular clock based on their estimate of 19.5% divergence from the crab *Eriocheir sinensis*, whereas we estimate distances at synonymous sites (*d*_s_) between *E. superba* and 13 other congeneric *Euphausia* species to be 27±7% (Table S7). This greatly exceeds the distance to the crab, suggesting their estimate may be downward-constrained by unusually conserved alignments. While we do not exclude the possibility of low or variable mutation rates among genomic regions and species, we find that comparable statistics for less constrained sites (e.g. *π*_S_ and *d*_s_) are not available, and call for additional studies of mutation rate in krill.

Other factors may also underlie low *N*_e_ / *N*_c_ ratios, such as highly skewed reproductive success among individuals, population size changes, linked selection that reduces variation around selected sites or constantly shifting adaptive peaks (Hedgecock and Pudovkin 2011; Charlesworth and Jensen 2022; Árnason et al. 2023), all of which suggest that natural populations may be far away from Hardy-Weinberg equilibrium. A recent study in the Atlantic cod found compelling genomic evidence of skewed reproductive success driven by pervasive and recurring selection (Árnason et al. 2023), evidenced for instance by negative Tajima’s D (*D*_T_) throughout the genome. We find strongly negative *D*_T_ in all krill and that genetic variation is much depleted at UTRs and non-synonymous sites, consistent with pervasive effects of purifying and linked selection across genomes. *Euphausia triacantha*, the least diverse species in terms of *π*_S_, has the largest excess of low frequency variants and the lowest *D*_T_ (−1.9), indicating recent population expansion in this species. Likewise, genome-wide patterns of variation in *E. superba* suggest it too has expanded from a recent bottleneck (Shao et al. 2023). Demographic history and natural selection together shape and limit variation in euphausiids.

### Genome scale assays provide new insights into euphausiid biodiversity

Our current understanding of marine zooplankton biodiversity is plagued by widespread cryptic variation, which has called for more systematic use of molecular tools in diversity assessments (Choquet et al. 2018; Bucklin et al. 2021; Bucklin et al. 2021). For instance, genome-wide data recently revealed the existence of three genetically distinct lineages in the pelagic pteropod *Limacina bulimoides*, originally considered a single circumglobal species (Choo et al. 2023). Because of their thin shells susceptible to dissolution, pteropods are used as indicators of ocean acidification. Thus, as different species may respond differently to acidification, an accurate definition of species boundaries is critical to assess its impacts.

In the krill, there are currently 86 described and recognized species, and no new species has been described since 1987 (Baker et al. 1990). Cryptic variation was reported within *Stylocheiron affine*, with a divergence level of 14% (mtCOI) differentiating Red Sea individuals from their Atlantic congeners (Wiebe et al. 2016; Bucklin et al. 2021). This mtCOI barcoding gap was on-par with interspecific levels of divergence in the genus (Bucklin et al. 2007). Here, we detect high mitochondrial divergence between *E. similis* and *E. similis* var. *armata*, that co-segregate with the main diagnostic trait. The *armata* variety was first described by Hansen (1911) as differing from *E. similis* by: ”[…] *a protruding, acute process on the third abdominal segment*”. Later, John (1936) added that “[…] *the process may be variable in size and shape; it varies from being low, rounded and inconspicuous to being a large compressed spine pointing backwards over the fourth segment*”, described slight differences in the structure of antennules and male copulatory organs, and proposed that *armata* reaches sexual maturity at a smaller length than *E. similis*. Russell et al. (1969) distinguished *armata* as living nearer the surface compared to *E. similis*. Our morphological analysis focused exclusively on the process on the third abdominal segment and we report a distinctive process in *armata* with a variable degree of development, in line with John’s findings (1936).

We estimate 6.2% of mitochondrial divergence (mtCOI) between *E. similis* and *armata*, above the within-species level of variation reported for the genus (average at 2.5%) (Bucklin et al. 2007), but below the between-species level (average 16.5%), making it inconclusive. However, the power of a single mitochondrial gene is limited. Here, we improve on this by also assessing variation across the genome. We found 36,930 fixed SNPs (with a *F*_ST_=1) across 4,011 genes. These are enriched for terms relating to roles in oocyte development and chromosome segregation (table S10), which could indicate barriers to gene flow (Hamaguchi and Sakaizumi 1992; Montecinos et al. 2017; Boynton et al. 2018). Our average estimate of genome-wide *F*_ST_ between *E. similis* and *armata* is 0.68, well above the *F*_ST_ of 0.41 reported between the two most differentiated lineages of the pteropod *Limacina bulimoides*, where Choo et al. (2023) concluded that these are most likely distinct species separated by strong reproductive isolation. Pairwise *dS* between *E. similis* and *E. similis* var. *armata* is 5.24%, slightly higher than between the two recognised species *E. spinifera* and *E. longirostris* (4%) (Table S7) although below the average between other sister species in the genus *Euphausia* (17%), which may suggest recent speciation. Considering all of the above, we propose to give the *armata* variety of *E. similis* the status of distinct species. Failure to recognize *E. similis* and *armata* as distinct evolutionary units or species would undercount krill biodiversity, inflate intraspecific structure and skew our understanding of species biology. In addition, it is essential to accurately describe taxonomy for our objectives of comprehending mechanisms underlying ecological adaptation in krill and assess adaptive potential to climate change.

### The genetic basis of ecological adaptation in krill

Polar regions and oceans are heating faster than the global average, impacting habitats and krill stocks in Arctic and Antarctic waters (Flores et al. 2012; Atkinson et al. 2019; Clem et al. 2020; Edwards et al. 2021; Rantanen et al. 2022). These rapid changes necessitate better insight into how thermal tolerance varies among species and what genetic mechanisms may contribute to adaptation to a warmer world. By analyzing CT50 data in six species, including new observations from subantarctic and temperate species, we find that CT50 thresholds appear to be correlated with the ambient temperatures of both sampling sites and habitats of species, likely reflecting adaptation to different thermal environments. However, when accounting for the different ambient starting temperatures, *E. crystallorophias*, *E. superba*, *E. vallentini*, *E. triacantha* and *T. inermis* show similar absolute CT50s, indicating comparable thermal tolerances beyond the temperatures experienced in their natural habitats. Only *M. norvegica* stands out with a much higher absolute CT50, which can be expected from its ubiquitous distribution. All the experiments carried out in this work were performed on animals collected at the colder end of their natural range, which may influence these patterns. By sampling each species also from warmer habitats, it would be possible to gain more comprehensive insight into their respective plasticities in thermal tolerance. Our observations suggest that these krill species may not be, at least at the adult stage, as thermally sensitive and stenothermic as other polar invertebrates (Peck et al. 2014). This is consistent with the findings of experiments conducted on *E. superba* (Toullec et al., 2020), but it may not apply during other stages of their life cycle, such as larval development, where growth rates and survival were shown to start deteriorating already around +3.0°C (Atkinson et al. 2006; Perry et al. 2020).

Daily and seasonal variation in ecological conditions such as light and temperature play critical roles in shaping the physiology and timing of behavior, growth, development and reproduction of zooplankton (Richardson 2008; Häfker et al. 2023), which in turn affect the function of marine ecosystems. The molecular mechanisms that govern important ecophysiological and phenological traits have only been intensely studied in some krill, focusing on species adapted to extreme photoperiodic seasonality at high latitudes (e.g. *E. superba*, *Thysanoessa* spp.) or that occur across wide geographical ranges (e.g. *M. norvegica*). This includes efforts to uncover and functionally characterize the photoreceptor and circadian clock gene repertoires (Biscontin et al. 2016; Christie et al. 2017; Palecanda et al. 2022; Urso et al. 2022) or genes involved in regulating the molting cycle (Seear et al. 2010), seasonal dynamics (Seear et al. 2012; Höring et al. 2021) and heat shock response (Huenerlage et al. 2016; Papot et al. 2016). However, analyses of how specific genes may contribute to environmental adaptation through natural selection have only been performed in heat shock proteins *hsp70* genes of Antarctic species before. Both *E. superba* and *E. crystallorophias* appear to have experienced diversifying selection between *hsp70* paralogs, but *E. superba hsps* are characterized by greater functional diversity and balancing selection associated with a more heterogeneous environment, while *E. crystallorophias hsps* may have evolved under stronger directional selection in a more constantly cold environment (Cascella et al. 2015; Papot et al. 2016). In the Antarctic krill, genes involved in controlling seasonal metabolic, molting, growth and reproductive cycles have been shown to be differentially expressed between summer and winter seasons (Höring et al. 2021; Urso et al. 2022). These cycles are thought to be driven by photoperiodic environmental cues such as day light, temperature and food availability and involve genetic pathways for photoreception and circadian clocks (Teschke et al. 2008; Biscontin et al. 2017; Piccolin et al. 2018; Höring et al. 2021). Environmental conditions vary widely among the habitats occupied by the species in our study and we find signatures of positive selection in hundreds of genes associated with multiple ecophysiological functions. We find selection signals across genes involved in photoreception (e.g. *ninaB*, *pdh* and *peropsin*), phenological physiology (e.g. *Eip75B*) and development (multiple cuticular proteins and chitinases), also in species evolving in warm environments.

While our candidate sets relate to diverse physiological functions, evolutionary change in thermosensory genes in particular could be key for adaptation to extreme or changing temperatures. In our analyses of Southern Ocean krill, we identify multiple candidate genes involved in detecting and responding to temperature, including a *TrpA1* homolog. TRP genes encode Transient receptor potential (TRP) Ca^2+^ ion channels, which are the main thermosensors in sensory neurons of animals (Dhaka et al. 2006; Bandell et al. 2007) and considered a “molecular toolkit for thermosensory adaptations” in animals (Hoffstaetter et al. 2018). TrpA1 is associated with harmful heat signaling and heat avoidance behaviors in both invertebrate and vertebrate ectotherms (Akashi 2021; Xiao and Xu 2021), as well as cold sensation in a wide range of animals (Akashi 2021; Zhang et al. 2022). In Southern Ocean notothenioid fish, *TrpA1* has been hypothesized to evolve through duplication and positive selection and be one of the main thermosensors underlying adaptation to cold Antarctic waters (York and Zakon 2022). In addition, we detect potentially adaptive substitutions across several genes encoding heat shock proteins or chaperonin-containing TCP1 (CCTs), which either assist with general or cytoskeletal protein folding. CCTs are eukaryotic “cold-shock” proteins and overexpressed during cold stress (Somer et al. 2002). In notothenioid fish, they appear to have undergone adaptive evolution to accomplish protein folding in cold and energy-depleted environments but are also upregulated from heat stress (Pucciarelli et al. 2006; Cuellar et al. 2014).

Our results suggest that homologous thermosensory receptors and protective proteins have been targeted by natural selection in both krill and fish, and that this repertoire of genes could underlie thermal adaptation also in other Antarctic or Arctic organisms. However, the specific functional properties of these candidate genes, and impact of their substitutions, are currently unknown in krill. For example, we detect signals of selection in a homolog of *subdued*. In *Drosophila*, the thermosensory Subdued ion channel collaborates with TRPs in heat nociception but activates at high temperatures that are unlikely to be experienced by any krill (+40°C) (Jang et al. 2015). On the other hand, *Drosophila* and krill Subdued homologs are only 43–45% identical and TMEM16 proteins overall are functionally diverse (Picollo et al. 2015). It is possible that krill Subdued requires considerably less activation energy, as seen in other cold-adapted proteins (D’Amico et al. 2002). The genes detected here are promising candidates but further functional validation is necessary to understand their role in adaptation.

### On adaptive potential in Antarctic krill

The potential for genetic adaptation ultimately depends on access to genetic variation, favorable demographics and life history traits. Therefore, the extent to which selection has shaped genetic variation among species in the past, and the potential they have to adapt to ongoing environmental change, may vary significantly among taxa. Empirical studies of protein evolution have indicated that few substitutions among apes have been fixed through positive selection (α=0–0.3) (Hvilsom et al. 2012; Galtier 2016), while high α in organisms like *Drosophila* (α≈0.5) and sea squirts (α≈0.8) indicate strong influence of positive selection (Smith and Eyre-Walker 2002; Tsagkogeorga et al. 2012). These analyses typically uncover higher α in invertebrates compared to vertebrates (Galtier 2016), find that adaptive evolution is limited by the supply of variation in low-*N*_e_ species and negatively correlated with “K-selected” life history strategies such as long lifespans or large offspring (i.e. the propagule size) (Galtier 2016; Rousselle et al. 2020), although how these potential determinants influence adaptation is still not well understood.

The hypothesis of high evolutionary potential in zooplankton (Peijnenburg and Goetze 2013) predicts that most variation between species should be shaped by adaptive evolution. This may not hold for krill, at least not judging by how natural selection has driven divergence in protein-coding genes in the past. Among the eight species of krill examined here, we find that the proportion of adaptive protein evolution (α) is only 0.40±0.11 (mean±SD), suggesting the majority of amino acid substitutions between species may have been fixed through processes other than adaptation. Compared to other arthropods, α seems low. For example, Galtier previously estimated α to 0.62±0.13 among 11 species (Galtier 2016). Our results indicate that the overall α in krill scales with *N*_e_, but it is more strongly influenced by non-adaptive change than by adaptive change. As seen in other species, the rate of fixation of slightly deleterious variants (for example due to drift in small populations) scales more clearly with *N*_e_ than the fixation rate of adaptive variants (Galtier 2016; Moutinho et al. 2020). Non-adaptive processes appear to contribute more to protein evolution in the Southern Ocean species *E. triacantha* (α=0.30) and *E. superba* (α=0.20). In the other krill, the ratios between *dN/dS* (ω) and *pN/pS* range from 1.29× to 1.83×, but are in these two species only 1.10× and 1.01×, respectively. For comparison, across the subset of 566 putatively cold-selected genes that we identify with branch-site tests and where these species have substitutions at selective sites, we find that α is 1.44× higher (0.43) in *E. triacantha* and 1.40× higher in *E. superba* (0.29) than the transcriptome-wide estimates, suggesting these genes have undergone more adaptive evolution overall than the genomic background. While the two species have undoubtedly adapted genetically to their respective environments, the question remains why so many substitutions throughout most genes have not evolved through adaptation. Our results suggest that K-strategists may be associated with low rates of adaptation in krill, as the two species have large bodies and larvae. However, demography and environment may also contribute. *E. triacantha* has the biggest excess of low frequency variants and lowest D_T_ (−1.9) in our analyses, consistent with recent expansion from a bottleneck of small *N*_e_ causing fixation of non-adaptive variants through genetic drift. Genome-wide analyses of *E. superba* suggest it too has recently undergone major population expansion (Shao et al. 2023), although our D_T_ is less extreme in this species (−1.4). An alternative explanation could be that slow rates of adaptation in the Antarctic krill may follow from comparably flat environmental gradients or slow rates of environmental change in Antarctic habitats. Since its formation, the Antarctic Circumpolar Current has acted as a barrier to warmer waters and helped maintain cold and stable Antarctic conditions (Clarke et al. 1992). Simulations indicate that the rate of recurring environmental change affects adaptive rates more strongly than *N*_e_ (Lourenço et al. 2013). Our sample is limited and only by expanding these analyses to small Antarctic species such as *E. crystallorophias* and *E. frigida* would it be possible to disentangle life history from habitat conditions as the main drivers of adaptive rate in Southern Ocean krill.

Krill are genetically adapted to their respective environments, but most divergence among species appears to have evolved through non-adaptive processes, suggesting adaptive traits have emerged over long timescales. What does this mean for their adaptive potential under rapid climate change? The answer lies in species-specific access to functional variation and efficacies of natural selection. We compared non-synonymous variation (*π*_N_) among our candidate genes for warm-adaptation against background *π*_N_ to gauge how much standing and assumably functional variation is available in populations to select from. Interestingly, we observe more standing variation for warm-adaptive genes among most species native to warm environments than to cold environments (supplementary fig. S12; Table S11). This could mean that more adaptive alleles are present in populations normally challenged by high or variable temperatures, possibly maintained by balancing selection or incomplete selective sweeps over environmental gradients or time. The Antarctic krill has no excess of functional (*π*_N_) variation among genes implied in warm-adaptation compared to background genes, and warm-adaptive genes have been evolving under even weaker influence of positive selection than the background (α=0.17 vs. α=0.20). A combination of long generation time (2–3 years) (Siegel 2000), extensive panmixia impairing local adaptation (Lenormand 2002; Shao et al. 2023) and possibly limited functional variation suggest the Antarctic krill has comparably low adaptive potential to rapidly changing environments.

## Conclusions

This study is the first to use comparative genomics at the genome scale to investigate evolutionary history and adaptation in marine zooplankton. We demonstrate the usefulness of transcriptome data as a means to identify patterns of genetic variation and adaptation among non-model species from different environments. Thorough understanding of taxonomy is essential to accurately interpret molecular signatures of adaptation in species and evaluate their adaptive potential. We identify significant genome-wide divergence between the two variants of *Euphausia similis* and *Euphausia similis* var. *armata* from the Indian Ocean. We show that the two forms are likely genetically isolated and evolving independently, with different levels of genetic diversity and rates of adaptive evolution. We argue that *Euphausia similis* var. *armata* should be given species status with the name “*Euphausia armata*”, resolving a 100-year old issue in krill systematics.

Conservation genetics has traditionally used neutral variation to survey population status and risk, but the link between neutral diversity and population viability is disputed and it can be argued that conservation should consider variation that directly promotes viability (Teixeira and Huber 2021). Comparison of new gene candidates for thermal adaptation, like *TrpA1*, *subdued*, *Hsc/Hsp70-interacting protein* (*HIP*), between tropical and Antarctic krill species may provide valuable information on functional variation that promotes fitness under warmer conditions. Surveys of genetic variation for these candidates can help uncover to what extent adaptive alleles occur in natural populations and provide insight into how much genetic change is required to adapt to future climates, i.e. the genetic offset (Fitzpatrick and Keller 2015).

We find that rates of adaptation are not uniformly high in krill, but vary widely among species. In particular, some Southern Ocean species like the Antarctic krill may have low adaptive potential. Just as other research indicates that there may be more structure and reproductive isolation among zooplankton populations than traditionally believed, our results suggest they do not behave like infinitely large effective populations unaffected by genetic drift or with unlimited adaptive potential. Instead, it seems to be determined by interactions between demography, life histories and environmental conditions. Expanding these analyses to more taxa could uncover common indicators of high or low adaptive potential in zooplankton, and allow more accurate forecasts of species distributions under continued change (Razgour et al. 2019; Capblancq et al. 2020).

## Materials and Methods

### Species identification and life histories

We used dissection microscopes and keys for morphological species identification, including Baker *et al*. (Baker et al. 1990), and *Cytochrome c oxidase I* (*COI*) barcodes for molecular species validation. Sample information is provided in Table S1. We georeferenced species ranges from (Brinton et al. 2000) on a OISST v2 Sea Surface Temperature (SST) map with *QGIS* http://www.qgis.org and computed the average SST for each range from color histograms in *GIMP* https://www.gimp.org/. To associate adaptive rates with life history traits, we compiled information from (John 1936; Russell et al. 1969; Baker et al. 1990; Brinton et al. 2000).

### Atlantic Ocean samples

We collected *M. norvegica* (Northern krill) specimens (n=7) from the Swedish Gullmarsfjord in 2018 using a small Isaacs-Kidd midwater trawl (IKMT) and preserved them in RNA*later*™ (Invitrogen™). We used material from three North Atlantic–Arctic *Thysanoessa* species: *T. longicaudata* and *T. raschii* specimens were collected in August 2012 in the Kongsfjorden (Spitzberg) and frozen in liquid nitrogen. We re-used a published RNA-seq library (ENA: SRR2174568) for *T. inermis* from (Huenerlage et al. 2016).

### Indian Ocean samples

Specimens of *Nematoscelis megalops* and six *Euphausia* species (*E. recurva*, *E. spinifera*, *E. similis*, *E. longirostris*, *E. triacantha*, *E. vallentini*) were collected in the Indian Ocean in 2019. We used a large IKMT aboard the R/V Marion Dufresne and kept specimens alive for 1–48 hours before preserving them in RNA*later*™ or performing experiments.

### Pacific Ocean samples

Four Pacific species of *Euphausia* were provided by Dr N. Tremblay: *E. distinguenda* and *E. lamelligera* (West coast of Mexico, February 2012), *E. pacifica* (West coast of Oregon, September 2011), *E. mucronata* (West coast of Chile, August 2011).

### Southern Ocean samples

During a 2017 REPCCOAI cruise, we collected a specimen of *T. macrura* North-East of Kerguelen Archipelago and a *E. frigida* specimen near the Indian Ocean Polar Front and froze them in liquid nitrogen. We downloaded a RNA-seq library for the Ice krill *E. crystallorophias* (ENA: ERR264582) from (Toullec et al. 2013) and 20 libraries for the Antarctic krill *E. superba* from (Höring et al. 2021) (NCBI: PRJEB30084).

### CT50 experiments

Experiments were performed using live *E. triacantha* and *E. vallentini* samples from the Indian Ocean and *M. norvegica* from the Arctic Ocean. The experimental protocol was identical to that used for *Euphausia crystallorophias* and *Euphausia superba* (Cascella et al. 2015) or the Arcto-boreal *Thysanoessa inermis* (Huenerlage et al. 2016). After acclimation of 24 h at ambient sea water temperatures, actively swimming animals were transferred to an experimental tank. Temperature was increased by 1°C per 10 min. The animals were maintained in the tank until they no longer responded to tactile stimuli of a probing rod. At this point, it was considered that the critical temperature had been reached. The CT50 was determined through the non-linear curve fitting option in Prism9 (GraphPad Software, LLC). The mobility/survival curve used was: Survival = c/(1 + (T/CT50), where c is the plateau value before the sharp decrease and CT50 is the threshold temperature at which only 50% of animals are mobile. The program explores the different parameter values and calculates a 95% confidence interval (standard error provided).

### Extraction and sequencing of RNA

For *M. norvegica* and samples collected from the Indian Ocean in 2019, we dissected and used abdominal muscle to extract RNA, using the Qiagen RNeasy® Plant Mini Kit and following the manufacturer’s protocol. Care was taken to remove the intestine to avoid bacterial or food contaminants. Thermo Scientific NanoDrop™ and Agilent Technologies 2200 TapeStation instruments were used to measure yield, purity and RIN-values. Samples with RIN>8 were provided to Science for Life Laboratory, Sweden, without DNase treatment (the kit protocol proved highly efficient in removing gDNA). Staff prepared Illumina RNA-seq 94 libraries using Illumina TruSeq Stranded mRNA Library Prep kit with polyA selection. Paired-end libraries (2×150 bp) were sequenced on a Illumina NovaSeq 6000 S4 lane and demultiplexed by the platform. Other samples were prepared as in (Huenerlage et al. 2016).

### Molecular species validation

For each library, we queried 1 million forward-oriented reads against all krill *COI* reference barcodes in the MetaZooGene database (MZGDB v3; n=3,003 sequences from 64 fully identified species) (Bucklin et al. 2021) using *BLASTN* (Tan et al. 2006; Camacho et al. 2009) and recorded best hits. For barcodes with 20+ hits, we computed the average identity scores. We took the barcode with the highest average score as an indication of species.

### RNA trimming, assembly and annotation

We used *Trim Galore!* v0.6.1 https://github.com/FelixKrueger/TrimGalore/ and *Cutadapt* v2.3 (Martin 2011) to trim low quality positions from reads (phred<20), only keeping trimmed reads of at least 50 bp. *Trinity* v2.11.0 (Grabherr et al. 2011) was then used with default settings to assemble 19 transcriptomes, making one “reference” transcriptome per species. When possible, we pooled reads from up to five specimens to maximize gene completeness. Preliminary tests indicated that pooling data produced on average 1.1–1.3× as many complete BUSCO genes compared to a large (44–133 M read pairs) or single small library (∼22 M read pairs). Due to unexpected population structure between *E. similis* and *E. similis* var. *armata*, a separate reference transcriptome was assembled for *armata*. The *T. raschii* transcriptome was assembled similarly to *T. inermis* in (Huenerlage et al. 2016). We used the Trinity script “*get_longest_isoform_seq_per_trinity_gene.pl*” to reduce redundancy by keeping only the longest splice isoform per gene.

We used *TransDecoder* v5.5.0 https://github.com/TransDecoder/TransDecoder to identify protein-coding transcripts and *TransDecoder.LongOrfs* to detect open reading frames (ORFs) >300 bp. ORFs were queried for domain homology against Swissprot with BLASTP v2.9.0+ (Camacho et al. 2009) (e-value cut-off 1e^-5^) and Pfam (release 34.0) with HMMER3 *hmmscan* v3.3 http://hmmer.org/. We used *TransDecoder.Predict* to identify and carry forward only protein-coding transcripts. Coordinates for functional regions (UTRs or coding sequence) were kept in GFF files. We used *KaKs_Calculator* (Zhang et al. 2006) to enumerate synonymous and non-synonymous sites, while transcriptome completeness was assessed using *BUSCO* v3.0.2b (Simão et al. 2015) and the arthropod odb9 lineage set from OrthoDB. We annotated the transcripts using queries against the *Drosophila* FlyBase database (dmel_r6.38) (Larkin et al. 2021) with *DIAMOND* v9.9.0 (Buchfink et al. 2015).

### SNP calling

We measured genetic variation within nine species/lineages: *M. norvegica*, *E. longirostris*, *E. similis* and its variant form *armata*, *E. spinifera*, *E. triacantha*, *E. vallentini, N. megalops* and *E. superba* using Single Nucleotide Polymorphisms (SNPs) detected in the RNA-seq data.

We mapped the trimmed RNA libraries from 113 specimens (3–20 specimens per species) to their respective reference transcriptomes using BWA-MEM v0.7.17-r1188 (Li 2013). Alignments were cleaned with *samtools* v1.14 (Danecek et al. 2021) to only retain reads mapping uniquely, concordantly and in pairs. Duplicates were removed using *Picard* v2.23.4 (http://broadinstitute.github.io/picard/). For each species, we called variants with *GATK* HaplotypeCaller v4.2.0.0. Variants were called per-individual and combined into a multi-sample gVCF with CombineGVCFs. Individuals were then jointly genotyped with GenotypeGVCFs. We used *vcftools* v0.1.16 (Danecek et al. 2011) to filter variants, retaining only bi-allelic SNPs genotyped by ≥5 reads at sites with a mean read depth of ≥5× across all individuals.

### Population structure

We here produced high-quality SNP subsets with vcftools, keeping SNPs with a minimum Phred quality score of 30, a minor allele count of three and less than 20% genotype missingness. We pruned SNPs based on linkage-disequilibrium using *Plink* v.1.90 www.cog-genomics.org/plink/1.9/ (Chang et al. 2015), proceeding in windows of 50 SNPs, sliding by 10 markers at a time and with a *r*^2^ threshold set at 0.8. We used the unlinked SNPs to investigate population structure in the Indian Ocean species by performing Principal Component Analyses (PCA) with Plink and ancestry analyses with *ADMIXTURE* v.1.3.0 (Alexander et al. 2009), using 2–N ancestral clusters (*K*) (N=number of geographic sampling sites).

### Divergence between E. similis (“similis”) and E. similis var. armata (“armata”)

Material from *similis* and *armata* was initially mapped to the same *E. similis* reference. In a joint SNP call, we uncovered extreme structure and used *vcftools* v0.1.16 to estimate the *F*_ST_ fixation index between them. We also observed that *similis* and *armata* samples matched different MetaZooGene *COI* barcodes. To further characterize this discrepancy, we assembled the *COI* mitochondrial gene (mtCOI) from the RNA-seq data. Reads with best hits against *E. similis* accessions (AF177186, MW210878 or MW210879) were extracted and assembled *de novo* into *COI* fragments using *SPAdes* v3.15.5 (Prjibelski et al. 2020). The fragments and the barcodes were aligned using *MAFFT* v7.453 (Katoh and Standley 2013). A pairwise distance matrix and neighbor-joining tree was inferred using *SplitsTree* v4.18.2 (Huson and Bryant 2006).

### Levels of intraspecific genetic variation

The full biallelic SNP datasets (above) were annotated with *SNPeff* v4.3T (Cingolani et al. 2012), i.e. to be synonymous, non-synonymous or UTR variants. We estimated variation using the population mutation rate Watterson’s theta (*θ*_w_) (Watterson 1975) separately at 5’-UTR, 3’-UTR, synonymous and non-synonymous sites, or jointly across full genes while accounting for accessible sites. We then used SplitsTree to calculate transcriptome–wide pairwise genetic distance matrices among individuals and deduce *π* (nucleotide diversity) and BioPerl to calculate Tajima’s D (D_T_) as an indicator of demographic history (Stajich et al. 2002). The effective population size (*N_e_*) was inferred from *N_e_* = *θ*_w_/4*μ*, where *μ* is the mutation rate per base and generation. Because mutation rates are unknown in krill, we used *μ* from snapping shrimp (Silliman et al. 2021), the closest species with a known rate.

### Orthology and phylogenetic analyses

We analyzed the 20 krill transcriptomes and seven outgroup datasets for gene orthology with *ProteinOrtho* v6.0.14 (Lechner et al. 2011), using *DIAMOND* to detect similarities. For each set of orthologous genes (an “orthogroup” or “OG”) with 10 or more krill species, we produced protein-level alignments. MAFFT was run using the G-INSI-I algorithm, using the “--allowshift --unalignlevel 0.8 --leavegappyregion” variable scoring matrix settings to reduce the risk of over-aligning non-homologous regions (Katoh and Standley 2016).

Single-copy OGs with at least 18 krill species and two outgroups were used to make a species tree. First, we trimmed unreliably aligned positions using *Gblocks* (Castresana 2000) (settings: “-t=p - b1=N -b2=N -b4=5 -b5=h -b6=y, where N was 50% + 1 of the number of sequences). We then performed phylogenetic inference of concatenated OGs using the maximum likelihood (ML) method in *IQ-TREE* v2.1.0 (Minh et al. 2020) and the JTTDCMut+F+I+G4 model (chosen using BIC) (Kalyaanamoorthy et al. 2017). We additionally produced individual gene trees using IQ-TREE (JTT+I+G4 model).

### Detecting positive selection and candidate genes for adaptation

We prepared OGs with 10+ krill species for analysis of adaptive evolution. The OGs were aligned without outgroups. For OGs with duplicate sequences (e.g. species-specific paralogs), we first produced gene trees with *Fasttree* (Price et al. 2010) and then used *OrthoSNAP* (Steenwyk et al. 2022) to split alignments into single-copy ortholog subsets, only keeping subsets with 10+ species. Nucleotide sequences were fitted to the protein alignments using *PAL2NAL* (Suyama et al. 2006). Spuriously aligned sequences may produce false-positive selection signals. We used *Gblocks* to trim unreliably aligned codons (settings: “-t=c -b5=h -b6=y”). In addition, we masked sequences around internal indels, replacing 4 codons with gaps around each indel and removing positions where >2 species had missing data. We also masked alignment fragments <15bp. For each OG, we pruned the species tree using *Phyutility* (Smith and Dunn 2008) to make unrooted species-matched subtrees.

We sought to uncover genes evolving under positive selection in krill adapted to either cold or warm climates and designed two tests. In the “cold” contrast, we took the terminal branches leading to five Antarctic–subantarctic *Euphausia* species as foreground branches, while assigning all other branches to the background. In the “warm” contrast, we took three Pacific tropical–subtropical equatorial/near-equatorial species as foreground branches. The ratio between non-synonymous substitutions per non-synonymous site (*dN*—a proxy for selection) over synonymous substitutions per synonymous site (*dS*—a proxy for unconstrained or neutral evolution), termed *dN/dS* or ω, compares the rate of amino acid substitutions between species against expectations under neutrality. Elevated ω (ω>1) can be used as an indication of positive selection, while ω=1 and ω<1 indicate absence of selection or purifying selection (Yang 2006), respectively. However, pairwise estimates for whole genes are only summary statistics that represent the long-term average mode of selection but do not localize selection to specific codons or lineages. Branch-site models leverage multiple sequence alignments and phylogenetic species trees and allow variable *dN/dS* ratios that can detect episodic positive selection involving only some branches (or species) and nucleotide sites (Yang and Nielsen 2002). We therefore performed branch-site tests with *codeml* in *PAML* v4.9j (Yang 2007) via the *ETE Toolkit* v3 (Huerta-Cepas et al. 2016), providing the codon-level alignment and subtree. We scanned for episodic positive selection by testing whether models allowing positive selection at one or more sites and branches fit the observed data and tree significantly better than null models that only allow neutral or purifying evolution (Yang and Nielsen 2002). For each OG, we ran *codeml/ETE* three times: i) using the bsA1 null model; ii) using the alternative bsA model that incorporates positive selection (Zhang et al. 2005); iii) performing a likelihood ratio test (LRT) to test if the null model could be rejected, taking the log-likelihood difference between models (−2ΔlnL) against a χ^2^ distribution (one df). For genes to be considered candidates, we required both a significant LRT in favor of the selective model and the detection of at least one positively selected site among foreground species, using Bayes Empirical Bayes posterior probability >95% (Yang et al. 2005).

To test for shared biological properties among the candidate genes, we performed Gene Ontology (GO) enrichment tests of *Drosophila* gene homologs using *ShinyGO* (Ge et al. 2020). We took the candidate genes as the target set and all analyzed genes as the background set.

### Estimating rates of adaptive evolution

McDonald & Kreitman introduced the idea to compare *dN/dS* (ω) between species against non-synonymous and synonymous polymorphisms segregating within species (*pN/pS*) to study selection at individual genes (McDonald and Kreitman 1991). Under the assumption that non-synonymous variants are either neutral, or deleterious and removed from populations via purifying selection, *pN/pS* should equal ω for genes evolving neutrally. Adaptive non-synonymous variants would be expected to be quickly fixed by positive selection and contribute to divergence (*dN*) but not polymorphism (*pN*), such that ω≫*pN/pS* can be taken as evidence for positive selection. By comparing ω to *pN/pS* across multiple genes, it is possible to estimate the overall proportion of adaptive amino acid substitutions (α), in which α = 1 - (*pN/pS* / ω), and infer how much of the observed variation between species may have been shaped by positive selection as opposed to neutral evolution (Smith and Eyre-Walker 2002). However, estimates of adaptive rates can be distorted by slightly deleterious non-synonymous variation that are not efficiently removed by purifying selection but segregate at low frequencies due to genetic drift, leading to overestimation of *pN/pS* and underestimation of *α* (Charlesworth and Eyre-Walker 2008; Moutinho et al. 2020), particularly when populations undergo bottlenecks. Current methods therefore interrogate whole site frequency spectra (SFS) of neutral and selective variants while aiming to control for confounding processes such as population structure, bottlenecks and interference between linked sites (Eyre-Walker and Keightley 2009; Messer and Petrov 2013; Galtier 2016; Al-Saffar and Hahn 2022). Therefore the site frequency spectrum (SFS) of synonymous variants is used to model demographic history, in order to more accurately model the distribution of fitness effects (DFE) from the non-synonymous SFS and derive more realistic estimates of *α* (reviewed in Galtier 2016; Moutinho et al. 2020; Al-Saffar and Hahn 2022), as well as the respective rates of adaptive or non-adaptive protein evolution (ω_A_ or ω_NA_).

For eight species with SNP data, we aimed to estimate the overall α, ω_A_ or ω_NA_ across thousands of genes. To estimate ω, we made 4-way datasets and pruned species trees that included the focal species and three additional species at different distances. We re-estimated branch lengths using the YN98+F1X4 model in *Bio++ bppML* v2.3.1 (Nielsen and Yang 1998; Guéguen et al. 2013), selected an ancestral node at appropriate distance and used *Bio++ bppancestor* v2.3.1 to infer the gene sequence at the ancestral node. The node was selected from overall pairwise genetic distances between species, avoiding very short distances as shared ancestral polymorphism may distort substitution-estimates (Mugal et al. 2020).

We then estimated synonymous/non-synonymous substitutions and sites between the focal and ancestral sequences using KaKs_Calculator and the “YN” method to account for unequal base frequencies, transition/transversion rate biases and multiple substitutions (Yang and Nielsen 2000). We compiled transcriptome-wide unfolded non-synonymous and synonymous site frequency spectra (SFS) from SNPs. We here used the ancestral sequence to polarize the SNP alleles: the allele shared with this sequence was considered ancestral, while the other was considered derived. This was performed with *basefinder* (a novel tool implemented here; commit adb820c) that cross-references SNP and alignment positions on either plus or minus strands. SFSs were produced from derived allele frequencies using genes with at least one polarized SNP. To have similar resolution of the SFSs between species and facilitate unbiased comparison, we downsampled the original population datasets to six or eight individuals. For each gene, individuals were selected to maximize the overall coverage and genotyped SNPs.

We used the model-based program *grapes* (Galtier 2016) to estimate α, ω_A_ and ω_NA_ in each species. The program infers fitness-effects and fits either non-parametric methods or models from *DoFE* (Eyre-Walker et al. 2006; Eyre-Walker and Keightley 2009) to the SFS + divergence data under maximum likelihood criteria. The “Basic” method derives the statistics by comparing *dN/dS* to *pN/pS* without correction, while “FWW” removes non-synonymous polymorphisms segregating at low frequencies (<15%) (Fay et al. 2001). In addition, *grapes* implements six model-based methods that estimate population demographics and fitness effects to correct for selectively neutral or fitness-altering non-synonymous polymorphisms (see Galtier 2016 and Al-Saffar and Hahn 2022 for details). For each species and method, we ran *grapes* 10 times with random initial values (“-nb_rand_start 10”).

## Supporting information

Comparative transcriptomics in krill Supplementary material

## Acknowledgements

This research was supported by grants 2018-04444 from the Swedish Research Council Vetenskapsrådet and the Inez Johanssons scholarship foundation awarded to AW. The computational infrastructure was enabled by project SNIC 2022/5-472 provided by the Swedish National Infrastructure for Computing (SNIC) at UPPMAX and the PDC Center for High Performance Computing, KTH Royal Institute of Technology, partially funded by the Swedish Research Council through grant agreement no. 2018-05973. We thank Dr N. Tremblay (Quebec university, Rimouski UQAR) for kindly providing Pacific Ocean krill samples, Camille Merland (Université Pierre et Marie Curie) and Marion Thellier (LOCEAN-IPSL, CNRS) for help with species identifications, the helpful research vessel and marine station staff that supported field sampling, Julien Yann Dutheil (Max Planck Institute for Evolutionary Biology) for helpful discussions and Mårten Larsson (Uppsala University) for folding the Peropsin protein. The REPCCOAI (Responses of the Pelagic Ecosystem to Climate Change in the Southern and South Indian Ocean) surveys were directed by Philippe Koubbi (Roscoff Biological Station) and J-Y. T. These surveys were supported by the French oceanographic fleet, the CNRS Antarctic Workshop Zone, the CNES KERTREND-SAT OSTST project led by Francesco d’Ovidio (LOCEAN), the European MESOPP H2020 programme and the TAAF National Nature Reserve programmes. We thank Philippe Koubbi for the opportunity to take part in these surveys.

## Author contributions

A. W. and J-Y. T. conceived and designed the study. A. W., J-Y. T., A. C., G. T., M. C., E. C. performed collection of data in the field, lab work or data curation. F. L. contributed software. A. W., M. C., J-Y. T. analyzed the data and wrote the manuscript. All authors reviewed and approved the final version of the manuscript.

## Data availability

The new sequence data underlying this article are available in the European Nucleotide Archive at https://www.ebi.ac.uk/ena/browser/home, and can be accessed with project accession PRJXXXXXXX. Annotated SNP datasets, transcripts and orthology datasets are available in the SciLifeLab Data Repository at XXXXXXXXXX.

## Code availability

Code for basefinder is available on Github: https://github.com/fellen31/basefinder.

## Notes

### Competing Interest Statement

The authors have declared no competing interest.

### Summary of Updates

* Addition of one co-author. * Minor revisions/clarifications of text. * Addition of author contributions and availability sections at the end. * Minor edits to supplementary materials.

