## Supplementary material for "Comparative population genomics provide new insight into the evolutionary history and adaptive potential of World Ocean krill": Comparative transcriptomics in krill Supplementary material

#### SUPPLEMENTARY TEXT

- Outgroup data for phylogenetic analyses.

#### SUPPLEMENTARY TABLES

Supplementary tables S1-S10 are provided as separate files.

- **Table S1.** Sample information for samples used in this study.
- **Table S2**. Characteristics of assembled reference transcriptomes.
- **Table S3**. Details of SNP datasets used for genome-scale analyses of genetic variation and population structure in nine krill species.
- **Table S4**. Characteristics of orthogroup sets used for phylogenetics or selection analyses.
- **Table S5**. Genes with signatures of positive selection.
- **Table S6**. Enriched gene ontologies among candidates genes.
- **Table S7**. Synonymous and non-synonymous divergence between species.
- **Table S8**. Rates of adaptive protein evolution.
- **Table S9**. Correlations between life history or ecological traits and rates of adaptive protein evolution.
- **Table S10**. Gene ontology enrichment among divergent genes between *E. similis* and *E. similis* var. *armata*.
- **Table S11**. Levels of non-synonymous variation in candidate genes for warm adaptation.

#### SUPPLEMENTARY FIGURES

Supplementary figures S1-S12 are provided in this document.

### SUPPLEMENTARY TEXT

#### Outgroup data for phylogenetic analyses

We used the peptide sequences from seven genome-sequenced crustaceans as outgroups in the phylogenetic analysis: the cladoceran *Daphnia magna* (Lee et al. 2019), copepod *Tigriopus californicus* (Barreto et al. 2018), amphipod *Hyallella azteca* (Poynton et al. 2018) and decapods *Penaeus monodon* (Uengwetwanit et al. 2020), *P. vannamei* (Zhang et al. 2019), *Cherax quadricarinatus* (Tan et al. 2020) and *Homarus americanus* (Polinski et al. 2021). Sequences for each species were downloaded from NCBI or genome portals.

### SUPPLEMENTARY FIGURES

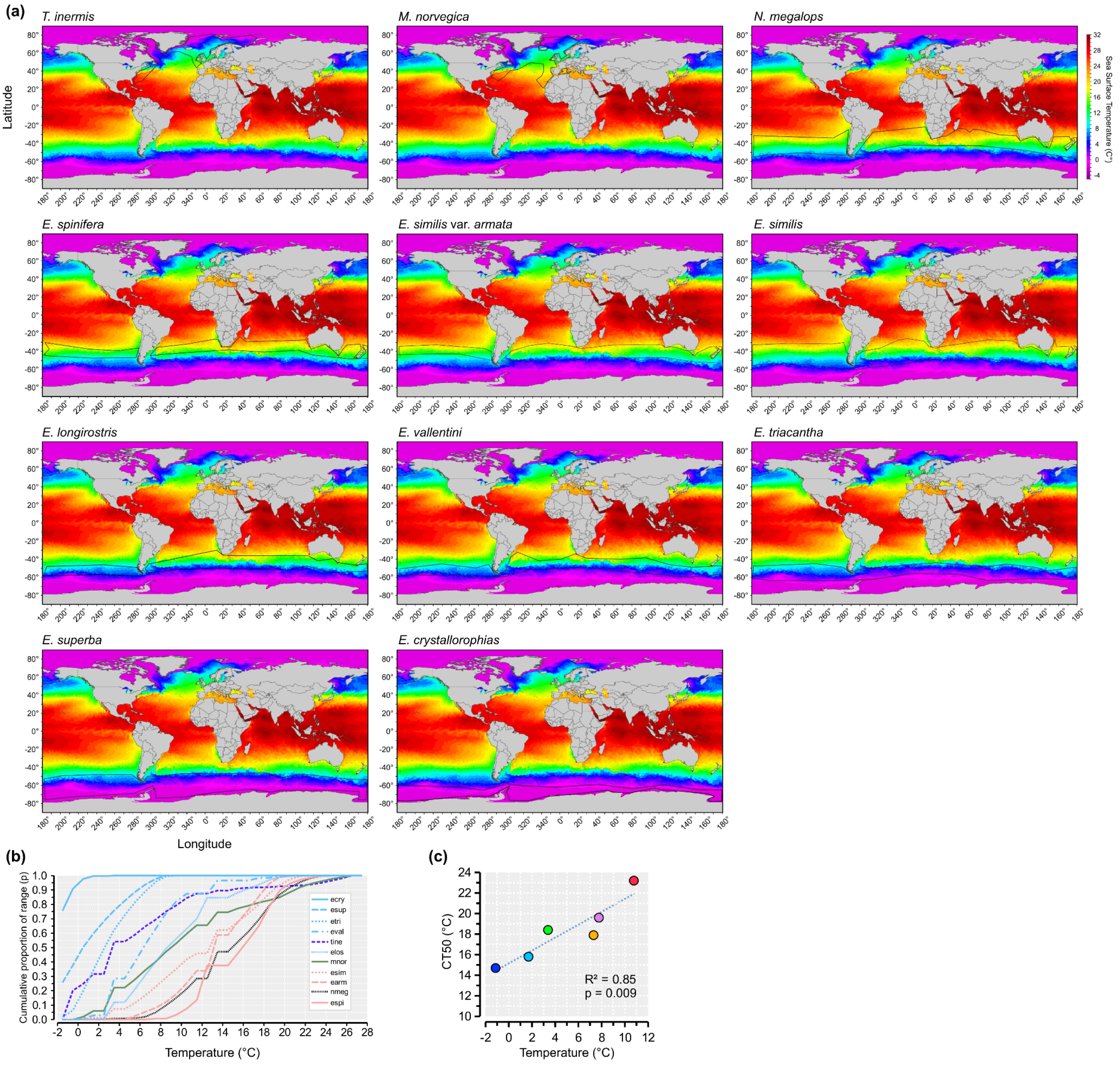


#### FIGURE S1

Ranges and thermal conditions of 11 krill species. (a) Species ranges (black lines) inferred from georeferencing distribution maps on top a map with OISST V2 Sea Surface Temperatures (SST) from ERDDAP (daily map snapshot 2022-06-01): https://www.ncei.noaa.gov/erddap/griddap/. (b) Temperature range of each species range shown as the cumulative proportion of 1°C intervals, sorted from low to high temperatures (ecry=*E. crystallorophias*; esup=*E. superba*; etri=*E. triacantha*; eval=*E. vallentini*; tine=*T. inermis*; elos=*E. longirostris*; mnor=*M. norvegica*; esim=*E. similis*; earm=*E. similis* var. *armata*; nmeg=*N. megalops*; espi=*E. spinifera*). (c) Linear regression correlation between CT50 and mean SST of the species range (same nine species as in fig. 2*B*).

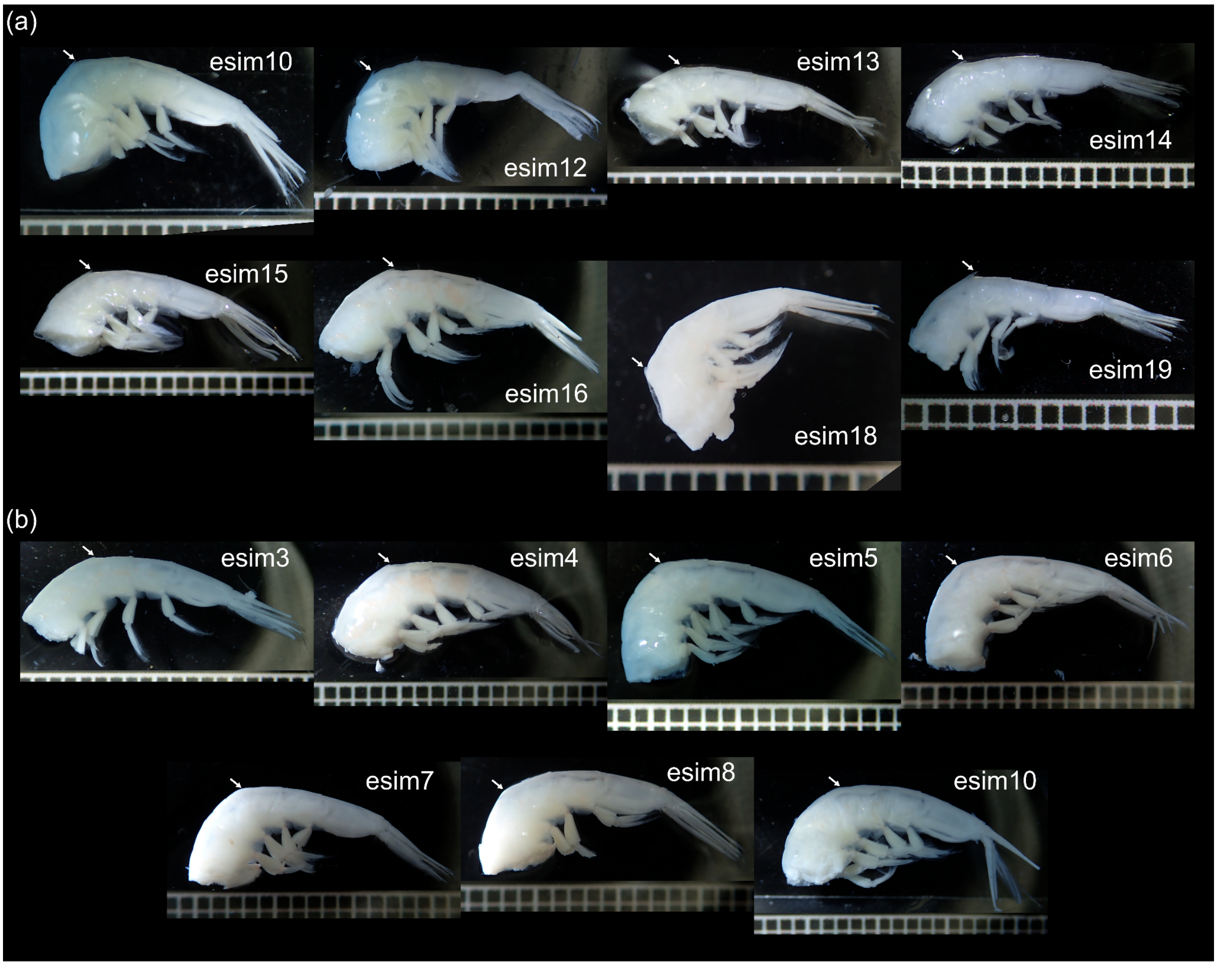


**FIGURE S2**

Photographs of the tails of *Euphausia similis* var. *armata* (a) (n=8) and *E. similis* (b) (n=7) samples that were collected from the Indian Ocean in 2019 and sequenced for this study. Tails were preserved in 95% EtOH. The arrow indicates the expected location of a dorsal accessory spine on the third abdominal segment. This spine is present in all *armata* samples (a) (although variable in size) but absent in all*similis* samples (b). Photos were taken with an Olympus TG-5 camera. A 1-mm grid is used for size reference. The *GNU Image Manipulation Program* was used to rotate photos, edit brightness and cut empty space between the mesh and sample. Additional samples that were identified in the field and sequenced in this study but are not shown here were preserved in RNA*later*™, and the abdominal segment was either consumed or poorly preserved for morphological analysis.

| | esim1 | esim2 | esim3 | esim4 | esim5 | esim6 | esim7 | esim8 | esim9 | esim11 | MW210878 | AF177186 | earm10 | earm12 | earm13 | earm14 | earm15 | earm16 | earm17 | earm18 | earm19 | MW210879 |
|---|---|---|---|---|---|---|---|---|---|---|---|---|---|---|---|---|---|---|---|---|---|---|
| esim1 | | | | | | | | | | | | | | | | | | | | | | |
| esim2 | 0 | | | | | | | | | | | | | | | | | | | | | |
| esim3 | 0 | 0 | | | | | | | | | | | | | | | | | | | | |
| esim4 | 0 | 0 | 0 | | | | | | | | | | | | | | | | | | | |
| esim5 | 0 | 0 | 0 | 0 | | | | | | | | | | | | | | | | | | |
| esim6 | 0 | 0 | 0 | 0 | 0 | | | | | | | | | | | | | | | | | |
| esim7 | 0.15 | 0.17 | 0.15 | 0.15 | 0.15 | 0.15 | | | | | | | | | | | | | | | | |
| esim8 | 0 | 0 | 0 | 0 | 0 | 0 | 0.15 | | | | | | | | | | | | | | | |
| esim9 | 0 | 0 | 0 | 0 | 0 | 0 | 0.15 | 0 | | | | | | | | | | | | | | |
| esim11 | 0 | 0 | 0 | 0 | 0 | 0 | 0.15 | 0 | 0 | | | | | | | | | | | | | |
| MW210878 | 0.23 | 0.24 | 0.23 | 0.23 | 0.23 | 0.23 | 0.45 | 0.23 | 0.23 | 0.23 | | | | | | | | | | | | |
| AF177186 | 4.85 | 5.07 | 4.85 | 4.85 | 4.85 | 4.85 | 5.02 | 4.85 | 4.85 | 4.85 | 4.73 | | | | | | | | | | | |
| earm10 | 6.25 | 6.6 | 6.25 | 6.25 | 6.25 | 6.25 | 6.4 | 6.25 | 6.25 | 6.25 | 5.86 | 7.28 | | | | | | | | | | |
| earm12 | 6.55 | 6.77 | 6.55 | 6.55 | 6.55 | 6.55 | 6.7 | 6.55 | 6.55 | 6.55 | 6.08 | 7.61 | 0.3 | | | | | | | | | |
| earm13 | 6.25 | 6.6 | 6.25 | 6.25 | 6.25 | 6.25 | 6.4 | 6.25 | 6.25 | 6.25 | 5.86 | 7.28 | 0 | 0.3 | | | | | | | | |
| earm14 | 6.58 | 6.77 | 6.58 | 6.58 | 6.58 | 6.58 | 6.73 | 6.58 | 6.58 | 6.58 | 6.08 | 7.64 | 0.3 | 0 | 0.3 | | | | | | | |
| earm15 | 6.18 | 6.53 | 6.18 | 6.18 | 6.18 | 6.18 | 6.33 | 6.18 | 6.18 | 6.18 | 5.63 | 7.12 | 0.3 | 0.6 | 0.3 | 0.61 | | | | | | |
| earm16 | 6.55 | 6.77 | 6.55 | 6.55 | 6.55 | 6.55 | 6.7 | 6.55 | 6.55 | 6.55 | 6.08 | 7.61 | 0.3 | 0 | 0.3 | 0 | 0.6 | | | | | |
| earm17 | 6.55 | 6.77 | 6.55 | 6.55 | 6.55 | 6.55 | 6.7 | 6.55 | 6.55 | 6.55 | 6.08 | 7.61 | 0.3 | 0 | 0.3 | 0 | 0.6 | 0 | | | | |
| earm18 | 6.55 | 6.77 | 6.55 | 6.55 | 6.55 | 6.55 | 6.7 | 6.55 | 6.55 | 6.55 | 6.08 | 7.61 | 0.3 | 0 | 0.3 | 0 | 0.6 | 0 | 0 | | | |
| earm19 | 6.1 | 6.44 | 6.1 | 6.1 | 6.1 | 6.1 | 6.25 | 6.1 | 6.1 | 6.1 | 5.63 | 7.12 | 0.3 | 0.6 | 0.3 | 0.6 | 0 | 0.6 | 0.6 | 0.6 | | |
| MW210879 | 5.88 | 6.08 | 5.88 | 5.88 | 5.88 | 5.88 | 6.1 | 5.88 | 5.88 | 5.88 | 5.86 | 6.97 | 0.65 | 0.22 | 0.65 | 0.22 | 1.09 | 0.22 | 0.22 | 0.22 | 1.09 | |

**FIGURE S3**

Uncorrected pairwise genetic distance matrix (% differences per bp) across the mitochondrial COI locus. Distances measured between *E. similis* ("esim") and *E. similis var. armata* ("arm") samples, as well as MetaZooGene reference sequences AF177186 (northern Pacific Ocean); MW210878 and MW210879 (the southern Atlantic Ocean).

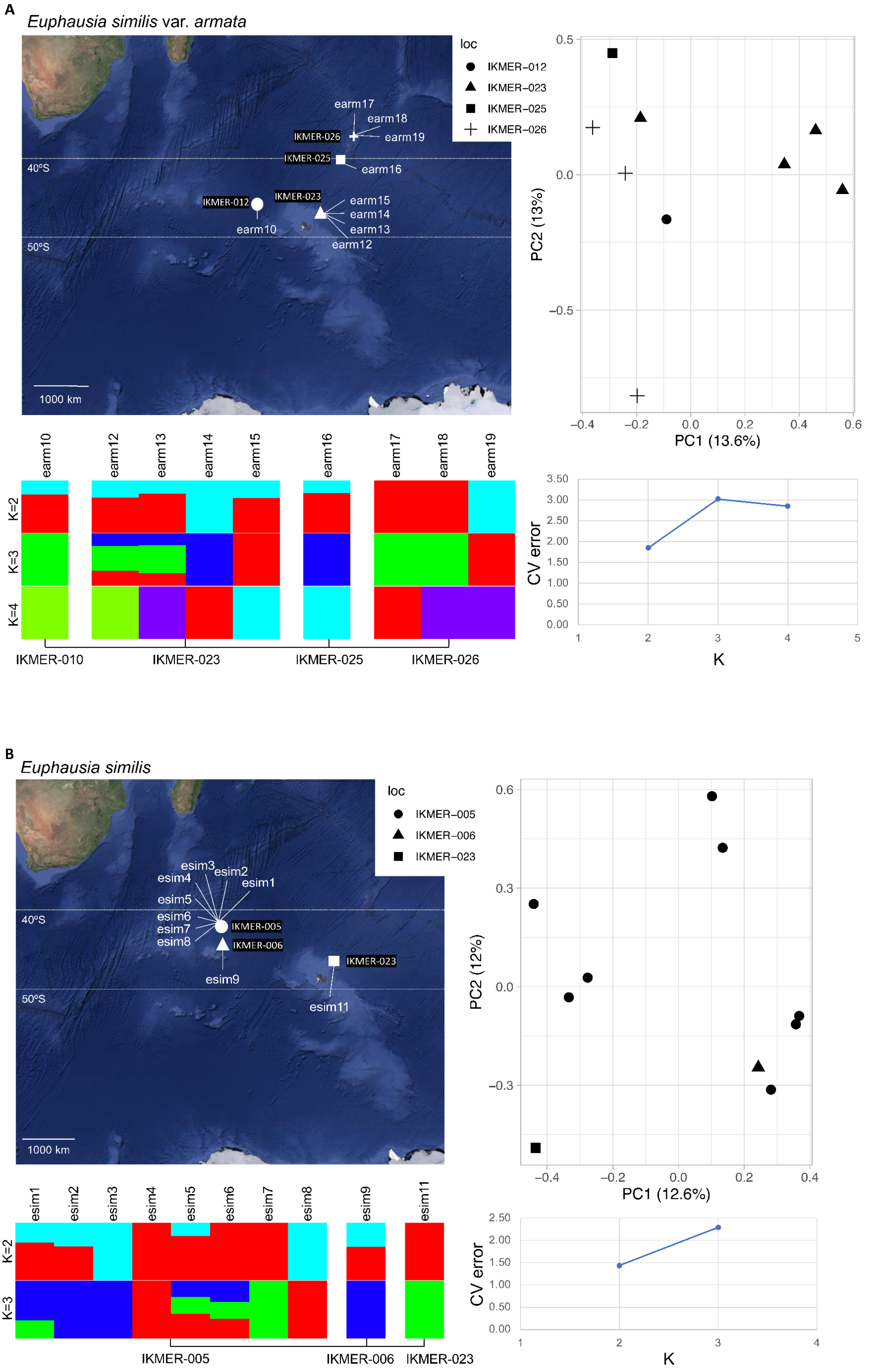


**FIGURE S4** (continued)

C

*Euphausia spinifera*

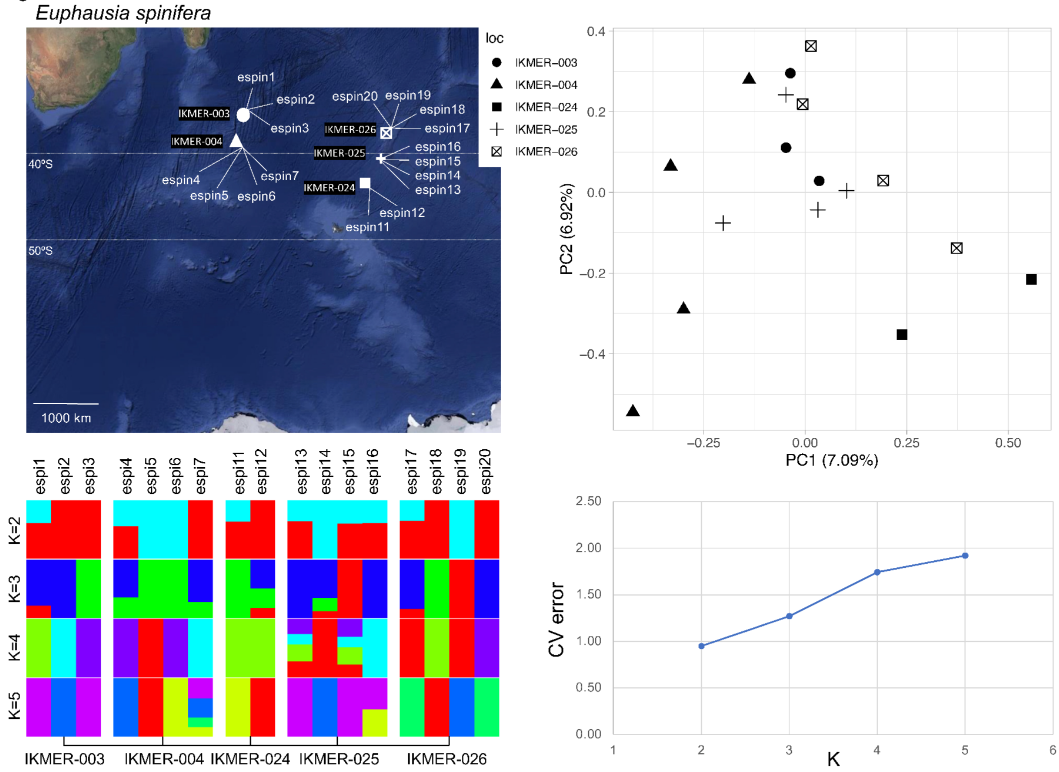


D

*Euphausia triacantha*

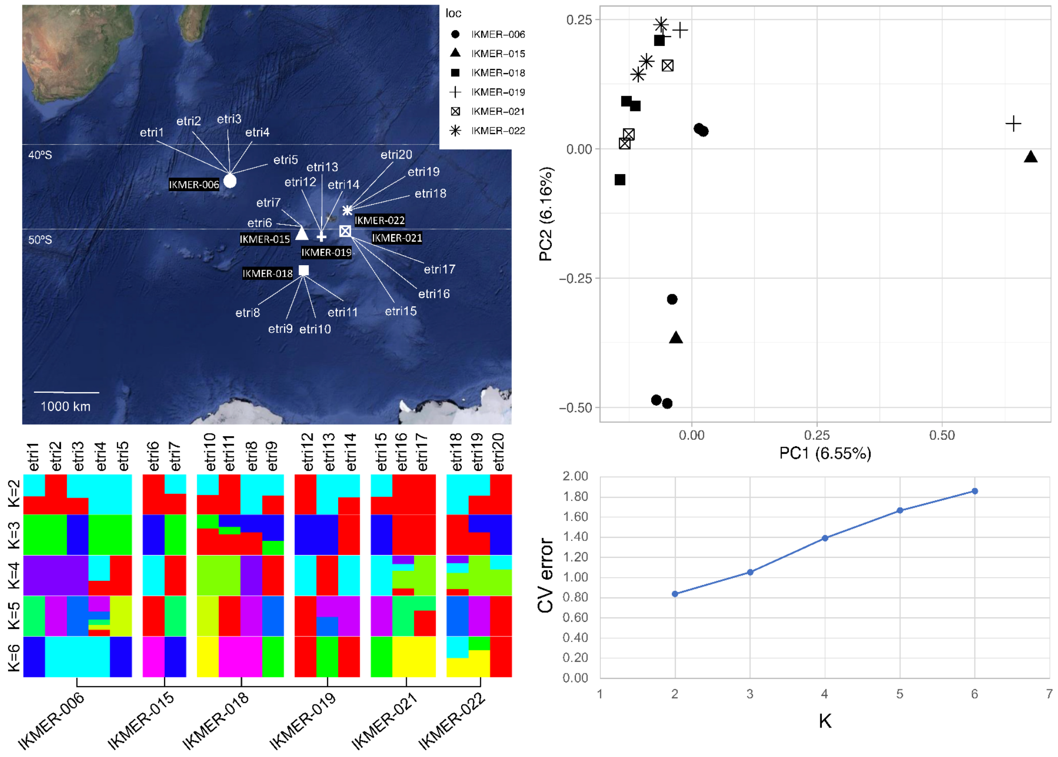


**FIGURE S4** (continued)

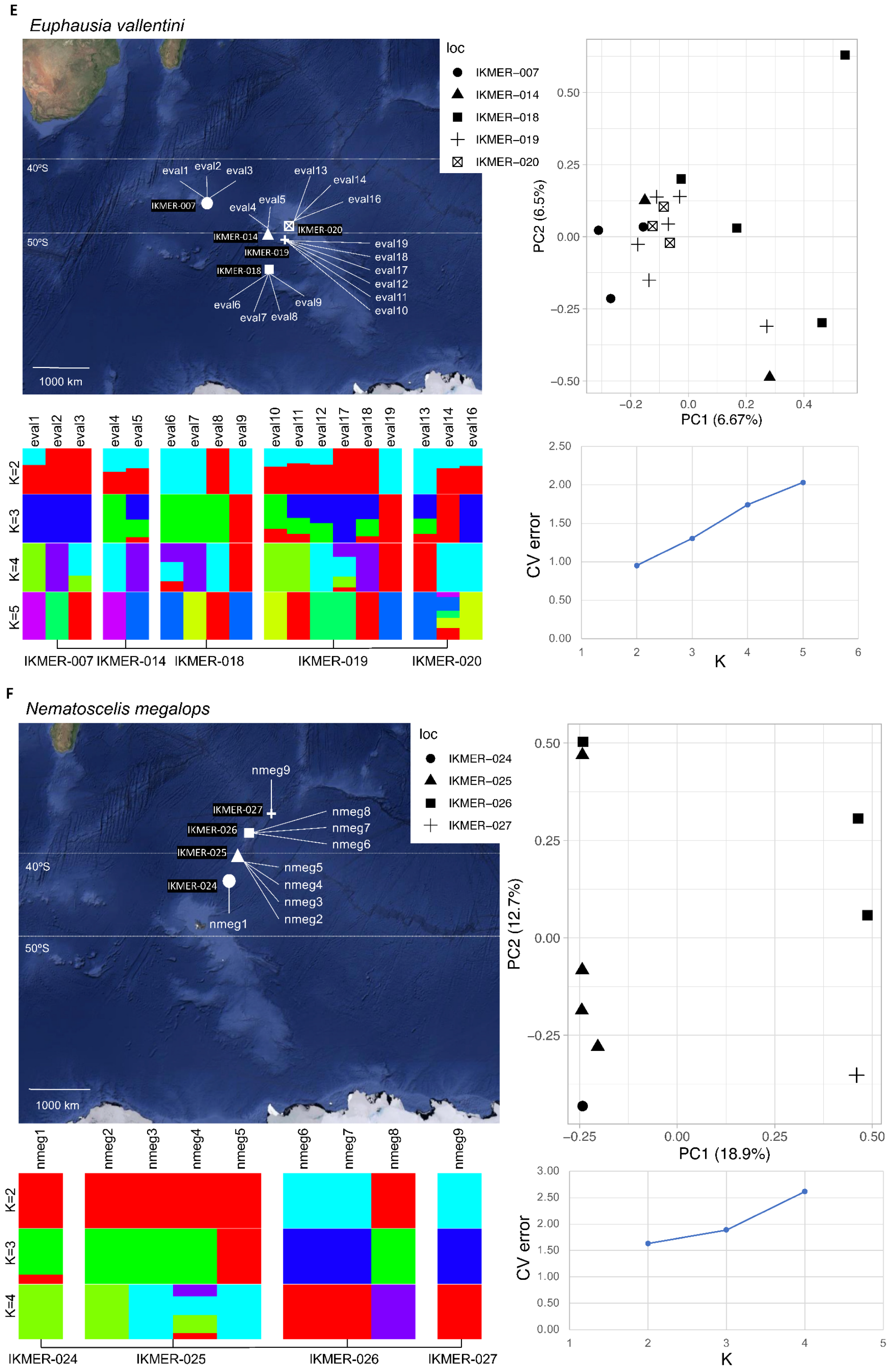


### FIGURE S4

Patterns of genetic population structure assessed from 32 k to 126 k independent SNPs per species in: **A**: *Euphausia armata*; **B**: *E. similis*; **C**: *E. spinifera*; **D**: *E. triacantha*; **E**: *E. vallentini*; **F**: *Nematoscelis megalops*. For each species, the top left panel shows a map of sampled locations and individual IDs; the top right panel is the representation of the two first components of a principal component analysis where each dot represents an individual from a location symbolized by a shape; the bottom left panel shows results of an ADMIXTURE ancestry analysis ran with K ranging from 2 to total number of sampling sites for that species; the bottom right panel shows the cross-validation (CV) error per each value of K. The value of K with the lowest level of CV error represents the most supported number of genetic clusters inferred by the analysis.

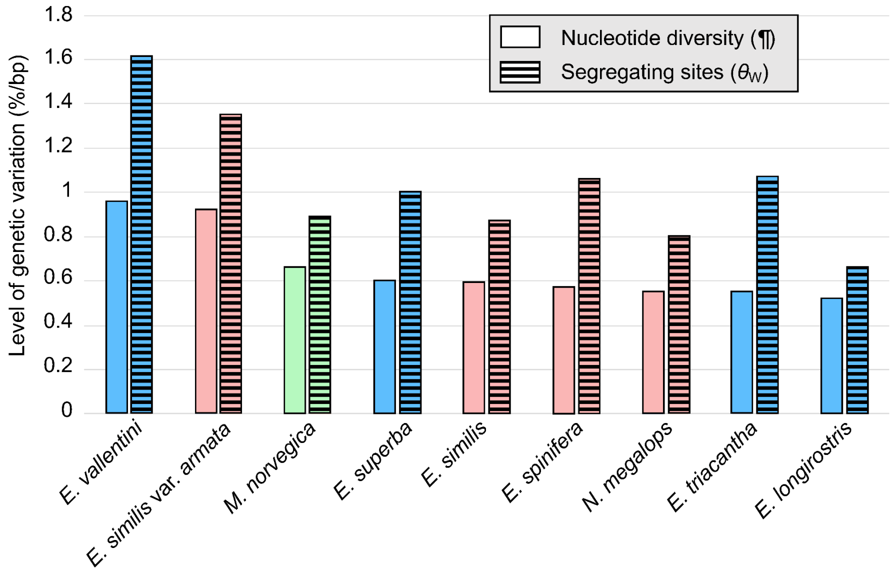


**FIGURE S5**

Transcriptome-wide levels of genetic variation in nine krill species. Nucleotide diversity ($\pi$) and Watterson's theta ($\theta_w$) were estimated from 259 k to 2 M SNPs per species, after correction for numbers of accessible sites.

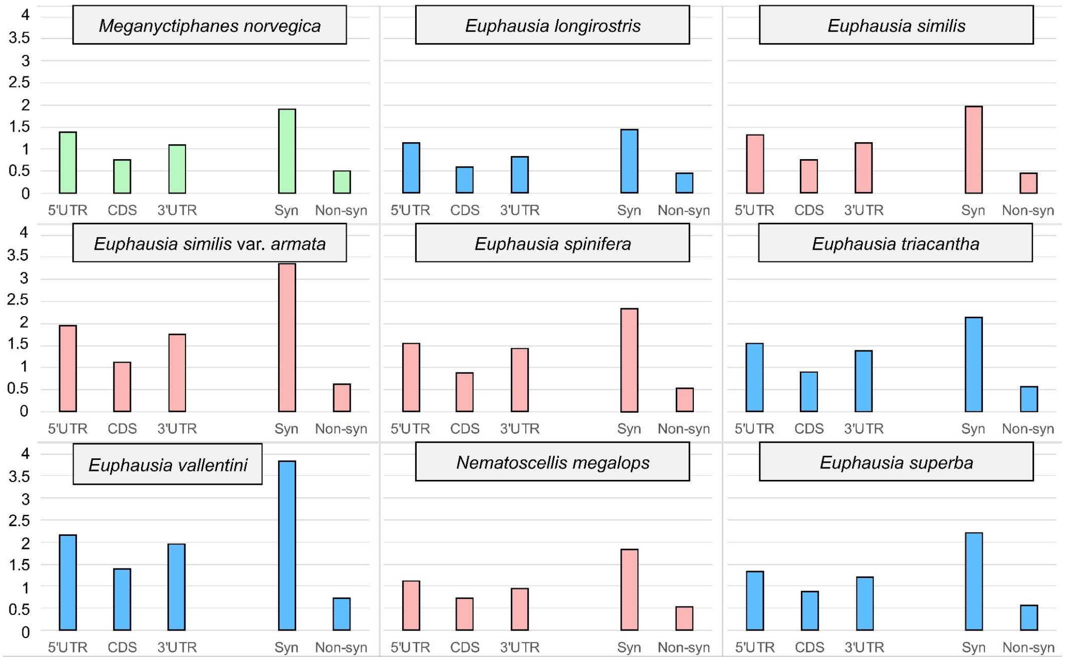


**FIGURE S6**

Average levels of genetic variation in nine krill species and five genomic regions. Variation was measured using Watterson's theta ($\theta_w$; % segregating sites per base) across untranslated regions (5'UTR and 3'UTR), coding sequence (CDS) and, more specifically, synonymous sites and non-synonymous sites within coding sequences. Color code for each species reflects the thermal habitat it is associated with (see fig. 1).

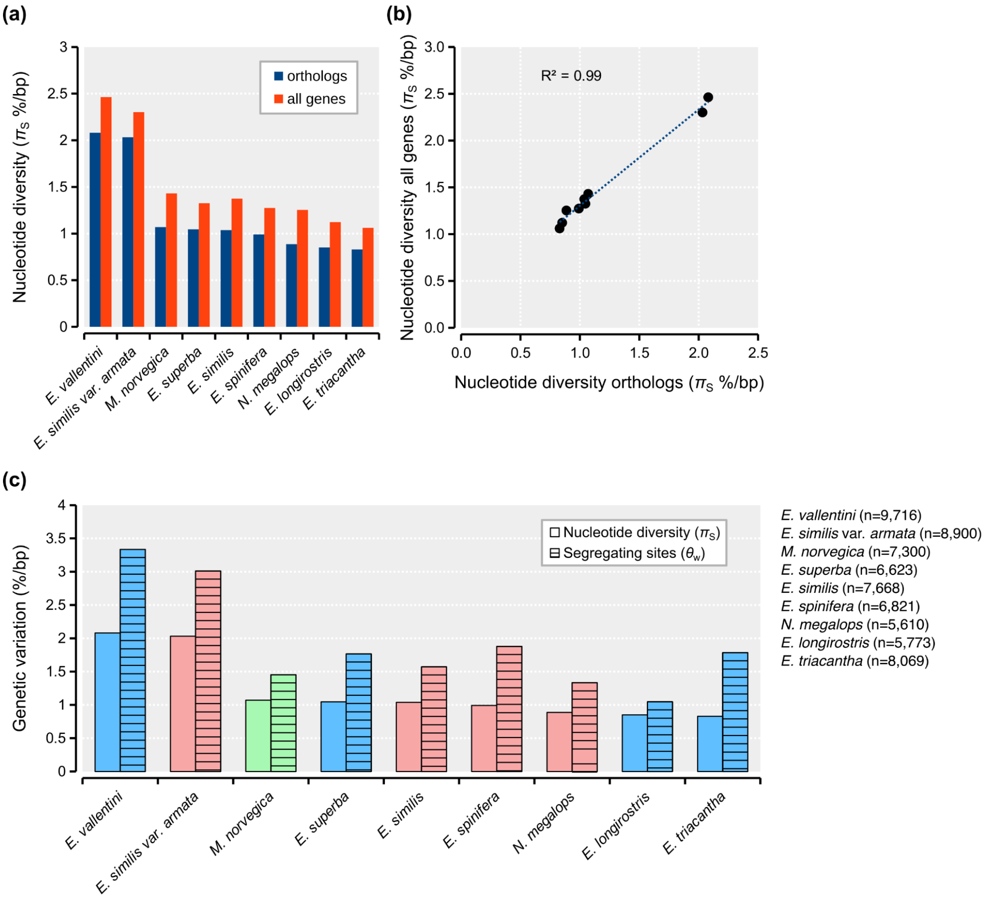


#### FIGURE S7

Levels of genetic variation at synonymous sites in a subset of n=13,255 orthologs (“orthologs”) with at least 10+ krill species compared against all polymorphic genes in each species (“all genes”). (a) Average nucleotide diversity across genes. (b) Linear regression between estimates, including the Pearson correlation coefficient. (c) Levels of genetic variation across the 13,255 orthologs measured using both nucleotide diversity and segregating sites (i.e. population mutation rate or Watterson’s theta). The number of genes used to measure each species is indicated (n).

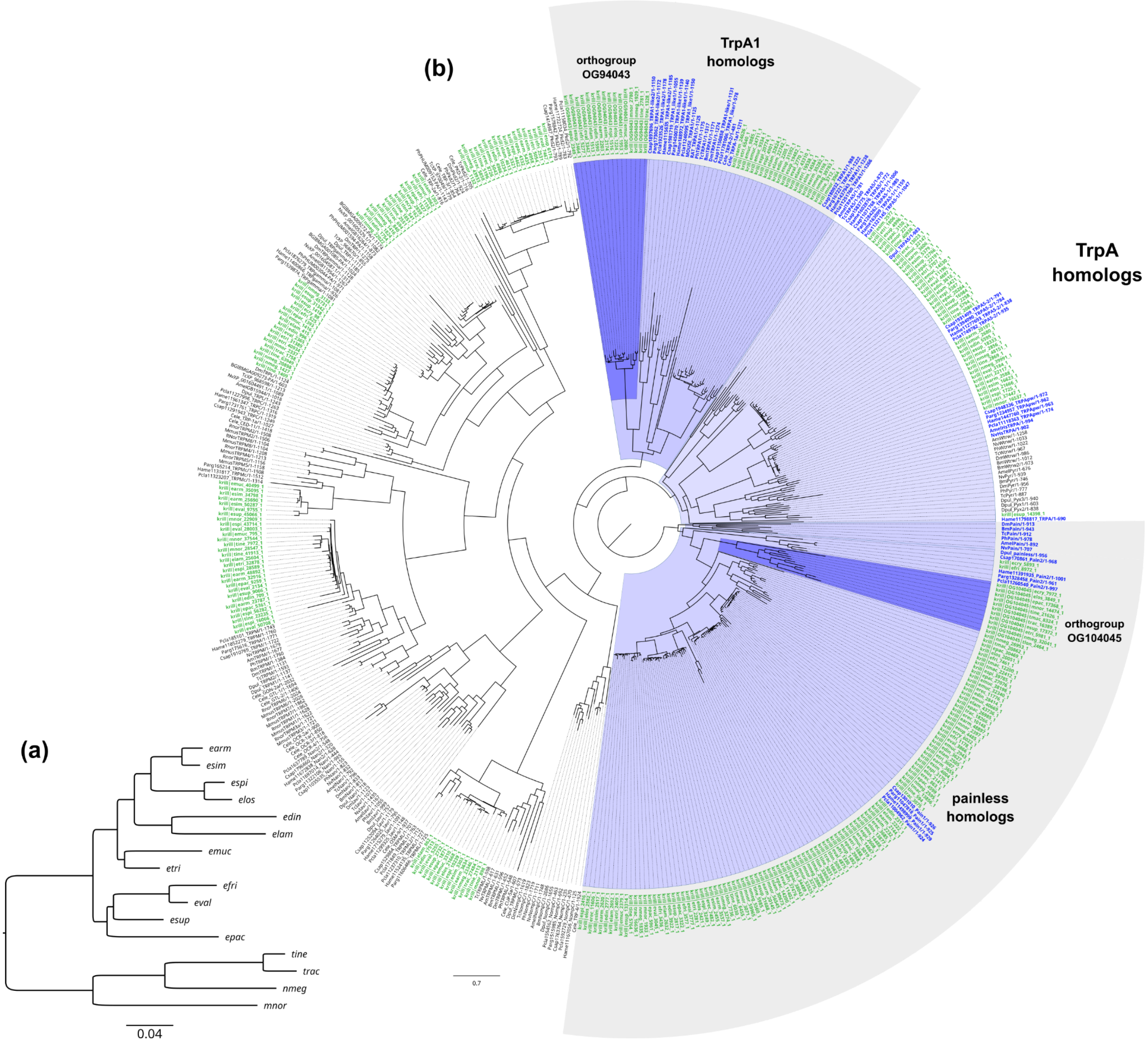


#### FIGURE S8

Phylogenetic assessment of two putative Transient receptor potential (TRP) homologs in krill: Transient receptor potential cation channel A1 (TrpA1) (orthogroup OG94043) and painless (orthogroup OG104045). Annotation using DIAMOND against *Drosophila* peptide sequences suggested OG94043 was a putative TrpA1 homolog, but with low identity scores (best hit: *Euphausia pacifica* "epac_2349_1" alignment against *Drosophila* FBgn0035934: identity=22.7%; length=480; e-value=4.2e$^{-16}$; score=85.5), while OG104045 was a painless homolog (best hit: *Thysanoessa longicaudata* "tlon_2464_1" alignment against *Drosophila* FBgn0060296: identity=26.1%; length=299; e-value=9.4e$^{-18}$; score=89.7). (a) Maximum likelihood tree for the OG94043/TrpA1 peptide sequences, indicating high consistency with the species tree. Tree generated with FastTree using the JTT model with CAT approximation with 20 rate categories. (b) Assessment of krill *TrpA1* and *painless* peptide homology towards insect and crustacean TRPs using the peptide dataset from (Kozma et al. 2020). We also queried the full krill transcriptomes against the TRP dataset using BLASTP to detect additional homologs and kept queries with identities >50% over 200 amino acids. Peptide sequences were aligned with MAFFT and a maximum likelihood gene tree was produced with FastTree as in (a). Krill sequences are indicated in green. Blue shade labels the branches corresponding to the TRPA subfamily and gray shades mark the TrpA1 and painless clades within this family. Crustacean TrpA1 and painless reference sequences are indicated in blue.

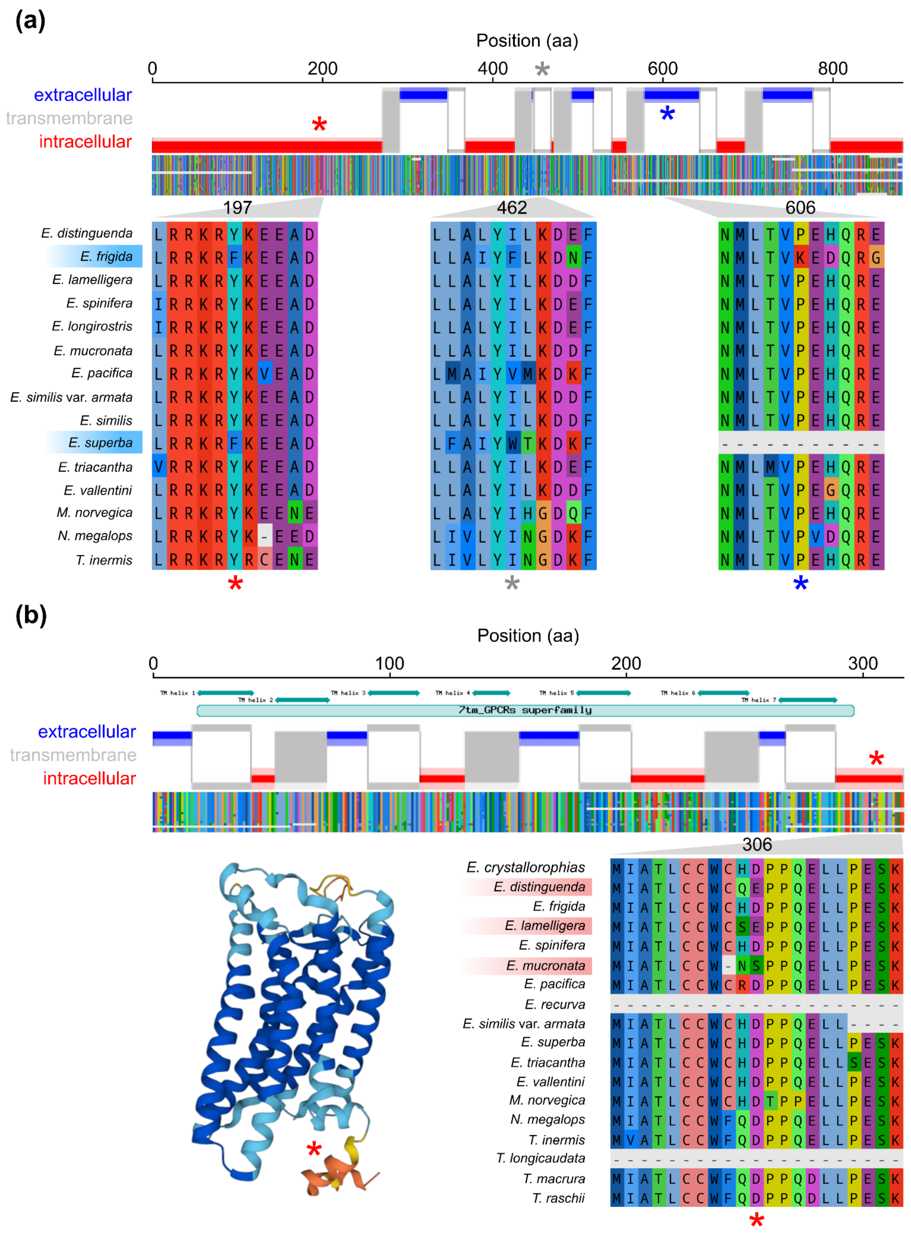


#### FIGURE S9

Examples of candidate genes for adaptation to cold or warm environments, detected using branch-site tests. (a) Signature of positive selection in the *subdued* transmembrane TMEM16 ion channel protein in cold-adapted krill. Top: the intracellular, extracellular and seven expected transmembrane domains of the protein encoded by the *E. valentini* ortholog (predicted using *TOPCONS v2.0*). Gray transmembrane regions=outbound; white=inbound. The three stars highlight putative positively selected sites in the protein (BEB>95%). Middle: the trimmed protein multiple-sequence alignment (MSA) of the krill orthogroup. Bottom: a magnified view of the MSA highlighting the site and species. (b) A signature of positive selection in the non-visual photoreceptor *Peropsin* gene in warm-adapted krill. Presentation layout as in (a). The intracellular, extracellular and seven expected transmembrane domains of the GPCR protein encoded by the *E. distinguenda* ortholog are shown (predicted using *TOPCONS v2.0*). Annotated domains were detected using BLAST Conserved Domain Search. The star indicates the putatively selected sites (BEB>99%). Bottom: a protein model produced with Alphafold and magnified view of the MSA highlighting the site and species.

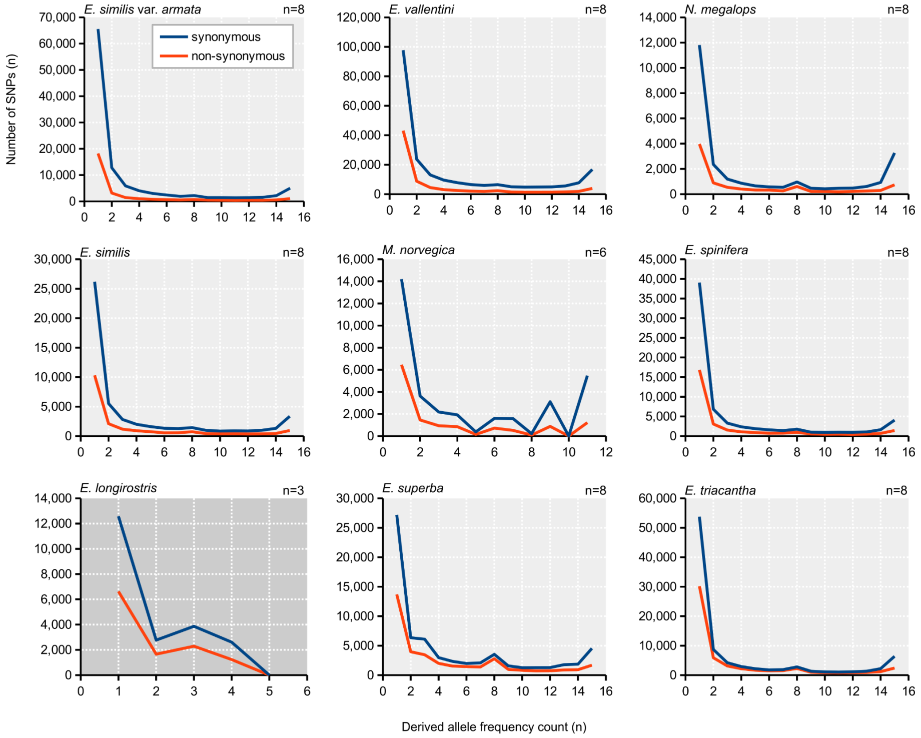


#### FIGURE S10

Unfolded site frequency spectra (SFS) for synonymous and non-synonymous variants detected in nine species of krill. Species names and sample sizes are indicated above each graph. Numbers of genes, SNPs and phylogenetic context used to polarize SNPs are indicated Table S8. *E. longirostris* was excluded from analyses due to small sample size and low resolution of the SFS.

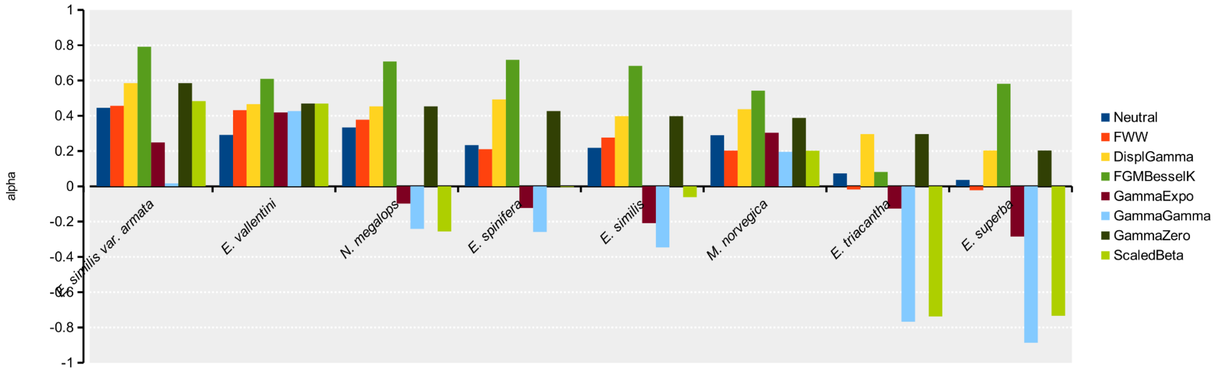


**FIGURE S11**

Proportions of adaptive protein evolution estimated using two basic non-parametric methods (Neutral and FWW) and six model-based methods (DisplGamma, FGMBesselK, GammaExpo, GammaGamma, GammaZero and ScaledBeta). Some models threw run-time errors or estimated nonsensical negative parameters for some datasets (e.g. ScaledBeta), suggesting poor fit between model and data.

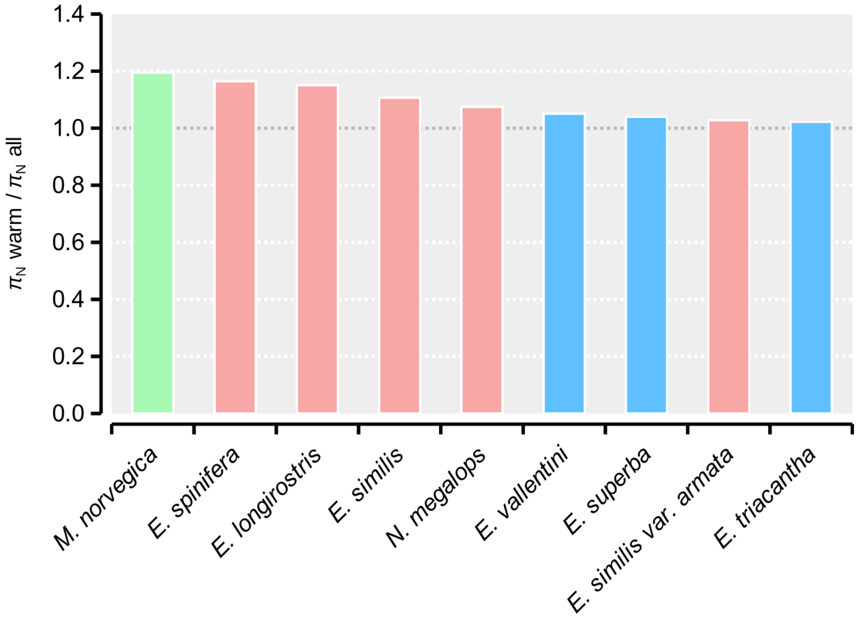


**FIGURE S12**

The ratio between non-synonymous nucleotide diversity measured across ~390 candidate genes for warm adaptation over non-synonymous nucleotide diversity measured for ~8,000 background genes. Details are available in Supplementary Table S10.
